# Individual humans attract different mosquito species

**DOI:** 10.64898/2026.01.30.702931

**Authors:** Kaylee M. Marrero, John S. Castillo, Dani Lucas-Barbosa, Anthony J. Bellantuono, Matthew A. Marrero, Dariel Cid, Andre L. Costa-da-Silva, Niels O. Verhulst, Matthew DeGennaro

## Abstract

Humans are not equally attractive to mosquitoes, leaving some more vulnerable to mosquito-borne illnesses than others. Body odor differences likely allow mosquitoes to discriminate between humans. Using a uniport olfactometer, we measured the attraction of *Aedes aegypti*, *Aedes albopictus*, and *Culex quinquefasciatus* mosquitoes for each of our 119 participants. *Ae. aegypti*, but not other species tested, were more attracted to male than female participants. Each of our three species ranked our participants differently, favoring a distinct subset of our cohort. For each species, mosquito attraction rates were used to define high and low attraction human odors and bacterial taxa. For example, *Ae. aegypti* and *Cx. quinquefasciatus* attraction was associated with the absence of odors like cyclic alcohols and monoterpenes, while *Ae. albopictus* attraction was associated with the presence of ketones. Each mosquito species exhibited distinct responses to individual humans, emphasizing both unique and shared cues for targeting their hosts.

## INTRODUCTION

Feeding on human blood is integral for the life cycle of highly anthropophilic mosquitoes such as *Aedes aegypti* and plays a significant role in species with a broader range of hosts like *Culex quinquefasciatus*. The behavior of *Ae. aegypti* and *Aedes albopictus* female mosquitoes spread flaviviruses that cause dengue, Zika, and yellow fever, while *Culex quinquefasciatus* can transmit the West Nile virus (1–3). In the Americas, all three mosquito species are predicted to increase their range putting new populations at risk (4). Mosquitoes use a multimodal approach to find a suitable host, integrating the detection of carbon dioxide (CO_2_), body odor, visual cues, humidity, and heat (5–12). Anthropophilic mosquitoes prefer urbanized environments and are highly attracted to human scent, which varies considerably from person to person (13–16). Mosquitoes have evolved to use odor to guide them to their preferred hosts (17–20). Still, human scent is a complex odor space comprising over 1,000 volatile organic compounds (VOCs), many uncharacterized, making it difficult to determine which odor plume components are responsible for mosquito attraction (21–25). Isolating the components of human odor that attract or repel mosquitoes could lead to novel strategies to combat vector-borne illness.

Human odor is largely derived from biotransformation of generally odorless skin secretions into VOCs by cutaneous bacteria of the human skin microbiome (26–29). The human skin microbiome is a dynamic, protective landscape of many different bacterial groups that utilize our sweat, sebum, and the stratum corneum as resources (30). Salient odors for mosquitoes may also be generated by UV light catalysis of sebum (31). Dominant taxa on our skin such as *Cutibacterium spp., Staphylococcus spp.,* and *Corynebacterium spp.* produce odors that contribute to an individual’s signature scent that can attract mosquitoes (32–35). Not only are the human skin microbiomes of individuals stable over time, but the diversity of species is correlated with health, signaling the importance of the immune system in maintaining healthy populations of skin bacteria (33). Mosquito attraction to humans is thus directly related to our skin microbiome, and individuals who are favored likely have a higher mosquito bite risk.

As humans encounter mosquito species with a range of anthropophilic behavior, it is important to understand the common cues that underlie their attraction as well as the specific cues that a mosquito species uses to find humans. Through the lens of single mosquito species, previous work has explored the connection between mosquito attraction rates across human hosts and their microbiome or volatilomes (36–41). It has become increasingly clear that there is a connection between human skin microbes and odor when mosquitoes target individual humans (28, 38–45). Cross-comparing attraction rates of multiple mosquito species to individual human volatilomes and skin microbiomes, to our knowledge, has not been previously accomplished. In this study, we compared mosquito attraction to 119 participants using three vector species found in the Americas. We found that *Ae. aegypti* were more attracted to males than females, but no sex bias was detected in *Ae. albopictus* or *Cx. quinquefasciatus*. We also found that different mosquito species are usually not attracted to the same participants. Odor and bacterial repertoires were associated with high attraction and low attraction participants for each species tested. The molecular signatures of mosquito attraction identified can inform next-generation repellent design and serve as indicators of mosquito bite risk.

## RESULTS

### *Ae. aegypti* show increased attraction for self-reported males over females

We recruited and tested 119 human participants from Miami, Florida. In addition to a demographically diverse group of volunteers, we studied the attraction rates of three different mosquito species, *Ae. aegypti, Ae. albopictus,* and *Cx. quinquefasciatus,* to these participants (Fig. 1A, Fig. S1). To measure this behavior for *Ae. aegypti* and *Ae. albopictus*, participants placed their left forearm inside a uniport olfactometer with female mosquitoes on the opposite side of the apparatus (46, 47). Since *Cx. quinquefasciatus* are night-biting mosquitoes, a nylon sleeve was used to capture the odors of each participant’s arms and experiments were conducted under nighttime conditions in the uniport olfactometer (39). We calculated relative attraction as the number of mosquitoes that entered the attraction trap divided by the total (n=30). *Ae. aegypti* showed an average attraction of 89% across all participants tested (Fig. S2A), whereas *Ae. albopictus* exhibited the lowest overall attraction of 27% (Fig. S2B). *Cx. quinquefasciatus* had an average attraction of 43% to human participant odor trapped on nylon sleeves (Fig. S2C).

**Figure 1.**
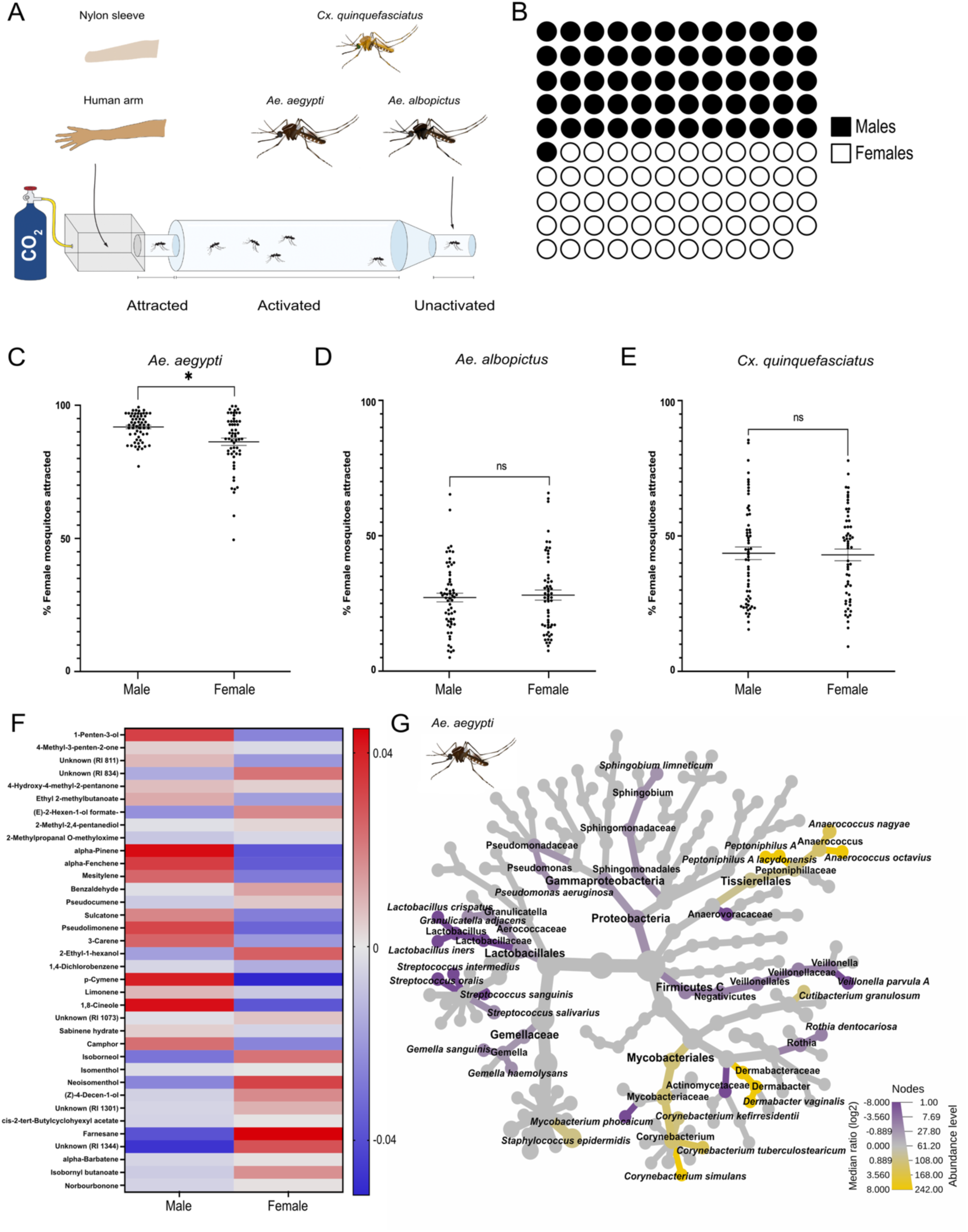
*Ae. aegypti* are more attracted to male than female participants. **(A)** Schematic of the uniport olfactometer used for quantification of mosquito attraction to individual human arms for *Ae. aegypti* and *Ae. albopictus* or humanworn nylon sleeves in *Cx. quinquefasciatus* experiments. **(B)** Distribution of self-reported sex for the participants (n = 61 males, 58 females). **(C-E)** Wilcoxon rank-sum test of mosquito attraction compared to all self-reported males and females in our study. **(F)** GC-MS heat map of chemical headspace for selfreported males (left) and females (right). Values listed represent the average chemical peak area. **(G)** Phylogenetic heat tree exhibiting a Wilcoxon rank-sum test of bacterial species abundance. Yellow nodes represent abundance in male participants, while purple nodes represent abundance in female participants. Labeled nodes represent taxa that are significantly abundant with an FDRadjusted p-value cutoff of 0.05.

To begin assessing attraction differences throughout the entire group, we analyzed the mosquito behavior, volatilome, and microbiome results of all self-reported male (n=61) and female participants (n=58) in our study (Fig. 1B). We compared the attraction rates for *Ae. aegypti, Ae. albopictus,* and *Cx. quinquefasciatus* and found that *Ae. aegypti* showed a significantly higher attraction for male participants (*P* =0.02) (Fig. 1C) whereas less anthropophilic species in our study did not show a difference in attraction between males and females (Fig. 1D & 1E). To ascertain the specific human odors that each mosquito species detects, the dynamic headspace from the left arm of each participant was collected. VOCs were identified by extracting volatiles from the left arm of participants, which was enclosed in a nylon bag for 90 minutes immediately after undergoing mosquito behavior experiments. Using gas chromatography-mass spectrometry (GC-MS), we analyzed the volatiles from the headspace of a human arm with k-means clustering into six different groups (Fig. S3). We found males exhibited a higher abundance of the alcohol, 1-penten-3-ol, and the terpenes, alpha-pinene, alpha-fenchene, p-cymene, limonene, and pseudolimonene, compared to females (Fig. 1F). For females, the terpene farnesane, the terpene derivative, isoborneol, and the cyclic alcohol, neoisomenthol, were more abundant on their skin compared to males (Fig. 1F).

Next, we compared the human skin microbiomes of males and females to evaluate the bacterial species that are associated with this difference in *Ae. aegypti* attraction. We sampled from the antecubital fossa and proximal volar forearm to explore the relationship of the arm with host-seeking variability in these vectors. We performed full-length 16S ribosome sequencing to identify bacteria comprising each participant’s human skin microbiome. Using a Wilcoxon rank-sum heat tree based on abundance level and median log ratio, we found several species that were significantly more abundant in males when compared with females, including *Anaerococcus nagyae*, *Anaerococcus octavius*, *Corynebacterium kefirresidentii*, *Corynebacterium simulans*, *Corynebacterium tuberculostearicum*, *Cutibacterium granulosum*, *Dermabacter vaginalis*, and *Staphylococcus epidermidis* (Wilcoxon test, FDR-adjusted *P* < 0.05) (Fig. 1G, Data S1). We found that two amplicon sequence variants (ASVs) of *C. acnes* were detected in all males, while only one ASV of *C. acnes* was detected in all females. The second ASV was detected in all female participants except one (Fig. S4). Although we detected sex differences in skin microbiota and odor profiles that correlate with *Ae. aegypti* attraction, further studies are required to determine if sex is an essential biological variable when this mosquito species targets an individual.

### Individual humans attract different mosquito species

Given the number of odors and bacteria that differ between our male and female participants, it is difficult to discern the salient cues for *Ae. aegypti* and impossible for *Ae. albopictus* and *Cx. quinquefasciatus*. To identify species-specific odor and bacterial signatures for mosquito attraction to humans, high and low attraction participants were separated for each species using a percentile analysis to isolate participants in the top 10% (90^th^ percentile) and bottom 10% (10^th^ percentile) of ranked mean attraction. This yielded 13 high attraction participants and 12 low attraction participants for *Ae. aegypti* (Fig. 2A & 2B), 12 high and low attraction participants for *Ae. albopictus* (Fig. 2C & 2D), and 12 high and low attraction participants for *Cx. quinquefasciatus* (Fig. 2E & 2F). A Wilcoxon test showed that there were significant differences in attraction between the high and low attraction groups for each species (Fig. 2B, 2D, & 2F).

**Figure 2.**
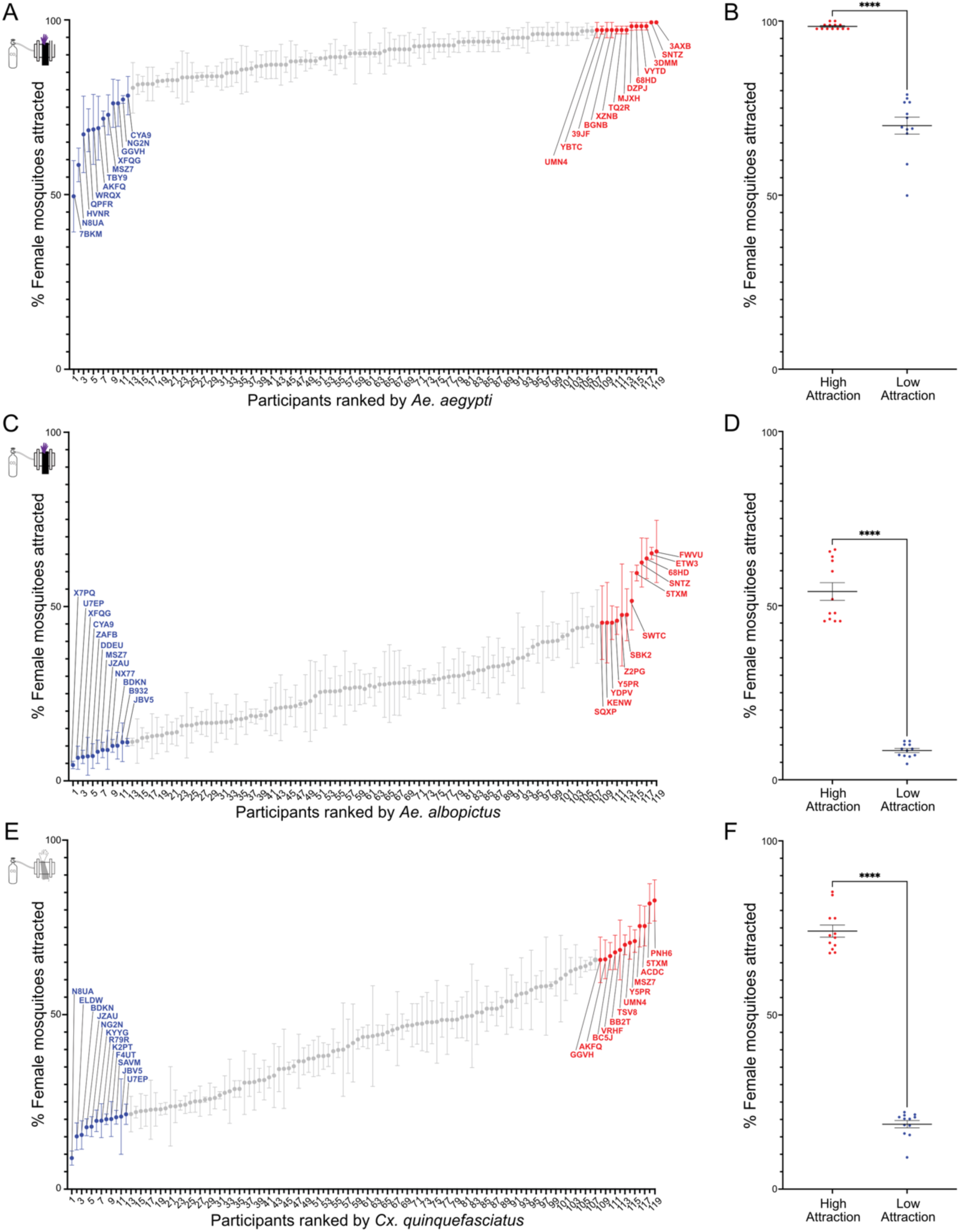
Distribution of participants by mosquito species attraction rates. **(A)** Attraction rates of female *Ae. aegypti* mosquitoes to individual humans. Participants are represented by a number ranked by ascending attraction to *Ae. aegypti*. Red dots reflect high attraction participants, while blue dots represent low attraction participants. **(B)** Wilcoxon rank-sum test comparing *Ae. aegypti* high and low-attraction groups. (p < 0.001). **(C)** Attraction rates of female *Ae. albopictus* mosquitoes to individual humans. Red dots reflect high attraction participants, while blue dots represent low attraction participants. **(D)** Wilcoxon rank-sum test comparing *Ae. albopictus* high and low attraction groups. (p < 0.001). **(E)** Attraction rates of female *Cx. quinquefasciatus* mosquitoes to human odor trapped on nylon sleeves. Red dots reflect high attraction participants, while blue dots represent low attraction participants. **(F)** Wilcoxon rank-sum test comparing *Cx. quinquefasciatus* high and low attraction groups. (p < 0.001). High and low attraction volunteer IDs are labeled.

We found that the participants that were most and least attractive for *Ae. aegypti* were often ranked differently by *Ae. albopictus* and *Cx. quinquefasciatus* (Fig. 3A). Overall, individual human attraction rates of *Ae. aegypti* and *Ae. albopictus* were weakly correlated (Pearson’s r=0.3492) (Fig. 3B). The correlation between individual human attraction rates of *Ae. aegypti* and *Cx. quinquefasciatus* was negligible (Pearson’s r=0.1431) (Fig. 3C). A similar result was observed for *Ae. albopictus* and *Cx. quinquefasciatus* (Pearson’s r=0.06034) (Fig. 3D). This suggests that each mosquito species is attracted to different individuals.

**Figure 3.**
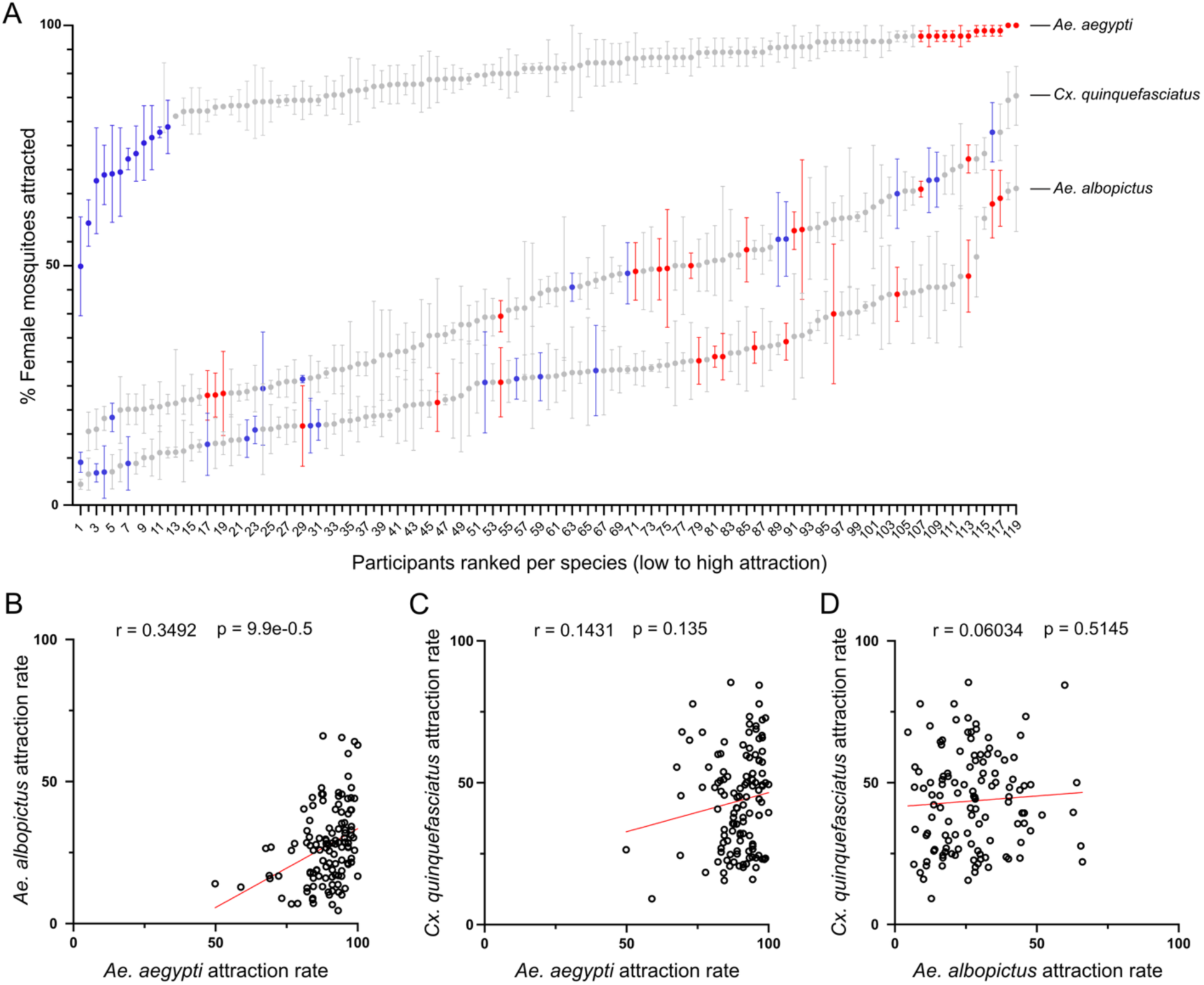
Individual humans attract different species of mosquitoes. **(A)** Behavioral responses in ascending order for all 119 participants to the three mosquito species studied. Red dots reflect *Ae. aegypti* high attraction participants, while blue dots represent *Ae. aegypti* low attraction participants. High attraction and low attraction groups were determined by percentile analysis for the greatest and lowest 10% of participants. **(B-D)** Pearson’s r correlation scatter plots of mosquito species ranked attraction to all participants. Each point represents each participant’s attraction rate for *Ae. aegypti* **(B-C)**, or *Ae. albopictus* **(D)** on the x axis and attraction rate for *Ae. albopictus* **(B)** or *Cx. quinquefasciatus* **(C-D)** on the y axis.

### Mosquito species are attracted by different VOCs from the human skin

Dynamic headspace analysis resulted in identification of VOCs associated with each mosquito species. We found only ethyl 2-methylbutanoate is more abundant in the high attraction group for *Ae. aegypti* than the low attraction group (Fig. 4A). Reduced attraction of *Ae. aegypti* was uniquely associated with higher levels of eight volatiles found in low attraction participants, with no species-specific odors associated with attraction identified (Table S1). The converse was true for *Ae. albopictus*; thirteen species-specific volatiles were more prevalent in high attraction participants than in low attraction participants. Among these was sulcatone, which has been linked to the evolution of mosquito preference for humans (18, 48). Only 2-ethyl-1-hexanol was species-specifically elevated in the *Ae. albopictus* low attraction group (Fig. 4B). Alpha-barbatene, alpha-fenchene, (E)-2-hexen-1-ol formate, (Z)-4-decen-1-ol, and pseudolimonene concentrations were elevated in participants that were highly attractive to *Cx. quinquefasciatus* (Fig. 4C). Limonene was more abundant in the headspace of low attraction participants for this species, while mesitylene abundance was the same in both high attraction and low attraction participants (Fig. 4C). Both compounds are worthy of further investigation as potential influencers of *Culex* species behavior. Interestingly, ethyl 2-methylbutanoate, (E)-2-hexen-1-ol formate, and alpha-pinene were all more prevalent in high attraction participants for all three vector species (Fig. 4D). Similarly, 1-penten-3-ol, 4-hydroxy-4-methyl-2-pentanone, 2-methylpropanal O-methyloxime, 3-carene, and isomenthol were all more prevalent in low attraction participants for all three species (Fig. 4E). Taken together, our results indicate that the presence of volatiles associated with low attraction in *Ae. aegypti* and *Cx. quinquefasciatus* may drive mosquito decisions to target individual humans. In contrast, *Ae. albopictus* may be guided to favored participants by attractive odor cues (Fig. 4D & 4E).

**Figure 4.**
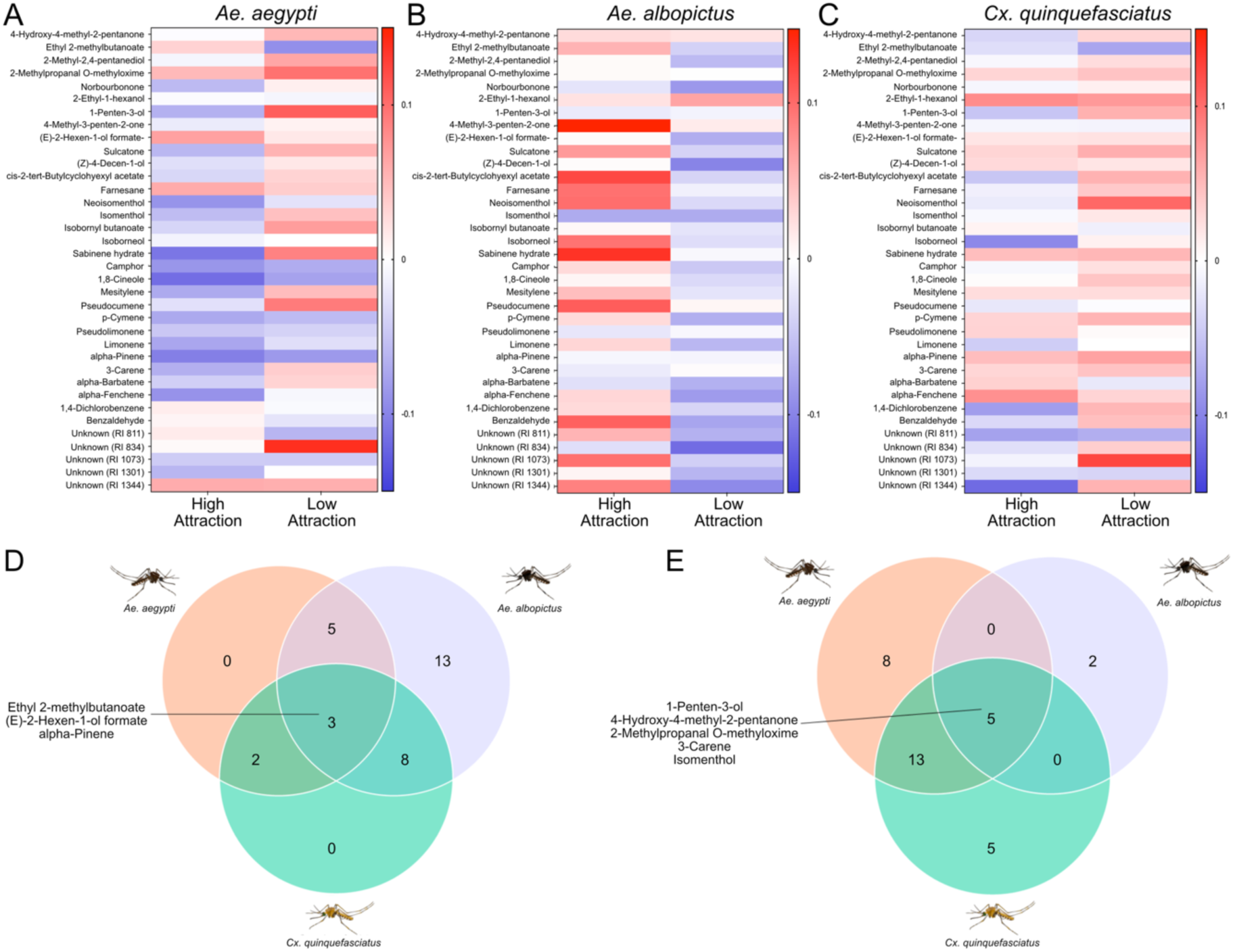
Distinct odor repertoires of individual humans are associated with species-specific mosquito attraction. **(A-C)** GC-MS heat map of chemical headspace for high (left) and low (right) attraction to **(A)** *Ae. aegypti,* **(B)** *Ae. albopictus,* or **(C)** *Cx. quinquefasciatus* mosquitoes. Values listed represent the average chemical peak area. **(D)** Venn diagram of compounds that are more prevalent in high attraction participants than low attraction for all mosquito species. **(E)** Venn diagram of compounds that are more prevalent in low attraction participants than high attraction for all mosquito species.

### Mosquito species are attracted to individuals with different skin microbial communities

We found each mosquito species’ attraction rates are associated with participants that possess distinct bacterial signatures. Examination of the skin microbiome for *Ae. aegypti* high and low attraction groups showed large amounts of *Staphylococcus* and *Corynebacterium spp.,* which are largely found across body sites (49, 50) (Fig. 5A). We found that the Firmicutes phylum, Bacili class, Staphylococcales order, Staphylococcaceae family, and *Staphylococcus* genus were enriched in participants that are highly attractive to *Ae. aegypti* mosquitoes. This is consistent with previous literature linking attraction of *Ae. aegypti* and *An. gambiae* mosquitoes to *Staphylococcus* bacteria (36, 38, 43, 51). For low attraction participants, *Pseudomonas A* and *Pseudomonas B* genera, which differ based on the structure of the flagella, as well as *Pseudomonas A stutzeri* and *Pseudomonas B oryzhibitans* were more abundant (52) (Wilcoxon test, FDR-adjusted *P* < 0.05) (Fig. 5B, Data S2). For sister species *Ae. albopictus*, we saw that the normalized read count was higher on average across the high attraction group than those of *Ae. aegypti* and *Cx. quinquefasciatus* (Fig. 5C, Fig. S5). We found *Sphingopyxis spp.*, *Sphingopyxis sp005503215*, *A. octavius*, *Rothia spp*., *Corynebacterium kroppenstedtii C*, and *Streptococcus intermedius* were significantly associated with the high attraction group whereas *Staphylococcus cohnii,* Anaerovaracaceae family, *Mogibacterium spp.,* and *Mogibacterium diversum* were significantly abundant in low attraction participants (Wilcoxon test, FDR-adjusted *P* < 0.05) (Fig. 5D, Data S3). Finally, for *Cx. quinquefasciatus* mosquitoes, we found that the normalized read count was lower on average for both the high and low attraction groups than those of *Ae. aegypti* and *Ae. albopictus* (Fig. 5E, Fig. S5). No bacteria were significantly associated with high attraction to *Cx. quinquefasciatus*, but *Actinomyces spp.* and *Actinomyces oris* were enriched in the low attraction group (Wilcoxon test, FDR-adjusted *P* < 0.05) (Fig. 5F, Data S4).

**Figure 5.**
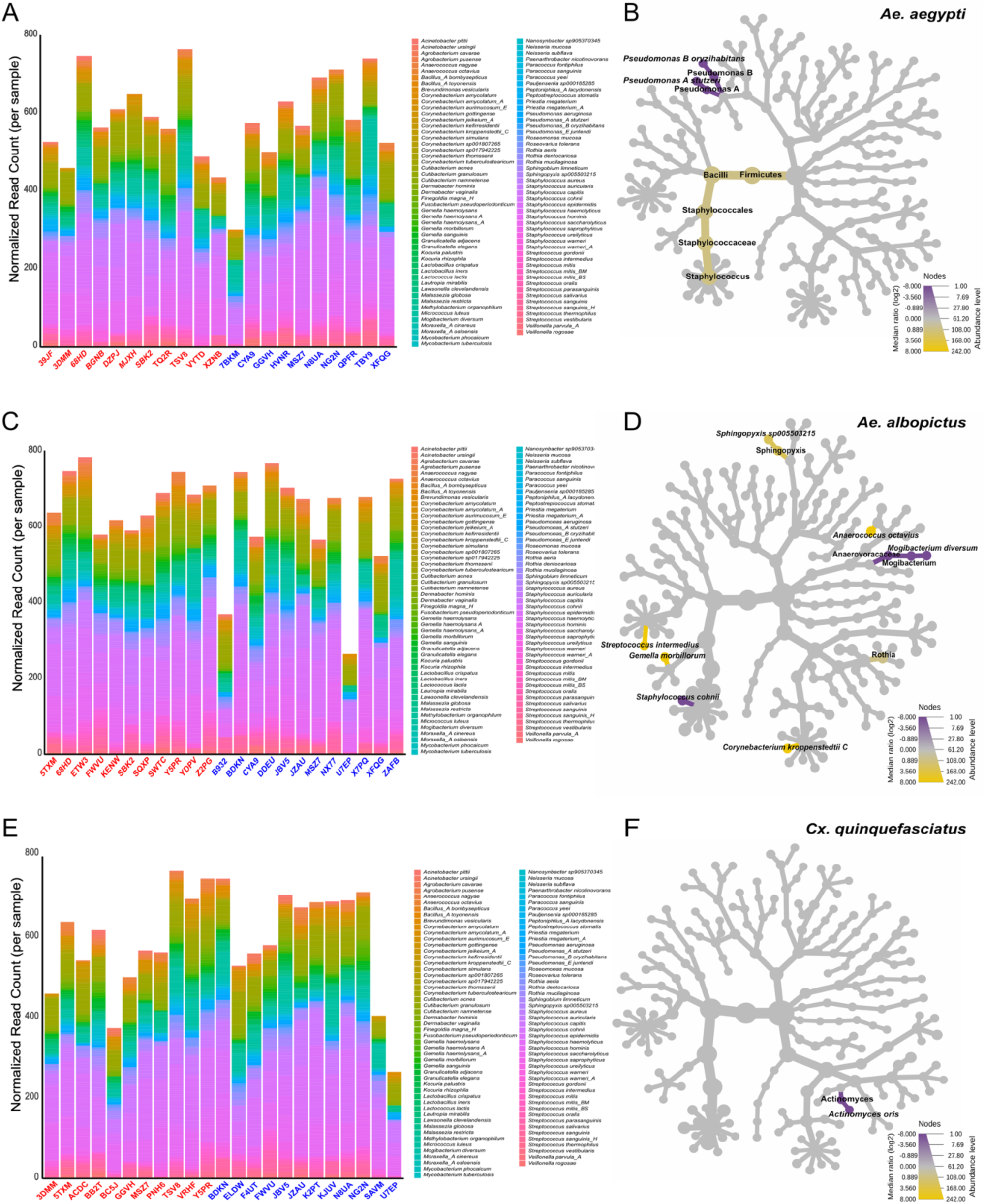
Mosquito species attraction rates are associated with distinct taxa of human skin bacteria. **(A, C, E)** Stacked bar chart depicting abundance of skin commensals on the arm of high and low attraction participants to **(A)** *Ae. aegypti,* **(C)** *Ae. albopictus,* or **(E)** *Cx. quinquefasciatus.* Bacteria are colored by species. **(B, D, F)** Phylogenetic heat tree exhibiting a Wilcoxon rank-sum test of bacterial species abundance. Yellow nodes represent abundance in high-attraction participants in **(B)** *Ae. aegypti,* **(D)** *Ae. albopictus,* or **(F)** *Cx. quinquefasciatus,* while purple nodes represent abundance in low attraction participants. Labeled nodes represent taxa that are significantly abundant with an FDR-adjusted p-value cutoff of 0.05.

### Distinct core human microbial communities correlate with species-specific attraction

We identified skin bacteria shared by participants in high or low attraction groups classified by mosquito species. Bacteria that were present for every participant of a high or low attraction group were assessed as the core microbiome for a given mosquito species. There were seven bacteria detected in all *Ae. aegypti* high attraction participants and eight bacteria detected in all *Ae. aegypti* low attraction participants (Fig. 6A). Some bacteria overlapped with the high attraction group, including both *C. acnes* variants, *C. granulosum,* and the same ASV of *S. epidermidis;* however, other bacteria were distinct for the low attraction core microbiome, such as *Micrococcus luteus* and *S. capitis.* Interestingly, *P. aeruginosa* was not detected in the high attraction core microbiome but was detected in the low attraction core microbiome. The core microbiome for *Ae. albopictus* high attraction participants had fourteen total bacteria and five in the low attraction group. The low attraction group also included the two ASVs of *C. acnes, Micrococcus luteus,* and *P. aeruginosa* (Fig. 6B). Notably, no *Staphylococcus spp*. appears in this low attraction core microbiome. Finally, thirteen bacteria strains were detected in all *Cx. quinquefasciatus* high attraction participants, including the two ASVs of *C. acnes, C. granulosum, P. aeruginosa,* three ASVs of *S. epidermidis,* and two variants of *S. hominis* (Fig. 6C). The low attraction core microbiome contained five species, including two ASVs of *C. acnes, S. capitis,* and one variant of *S. hominis*. This was the only group to contain *Corynebacterium tuberculostearicum* in any core microbiomes.

**Figure 6.**
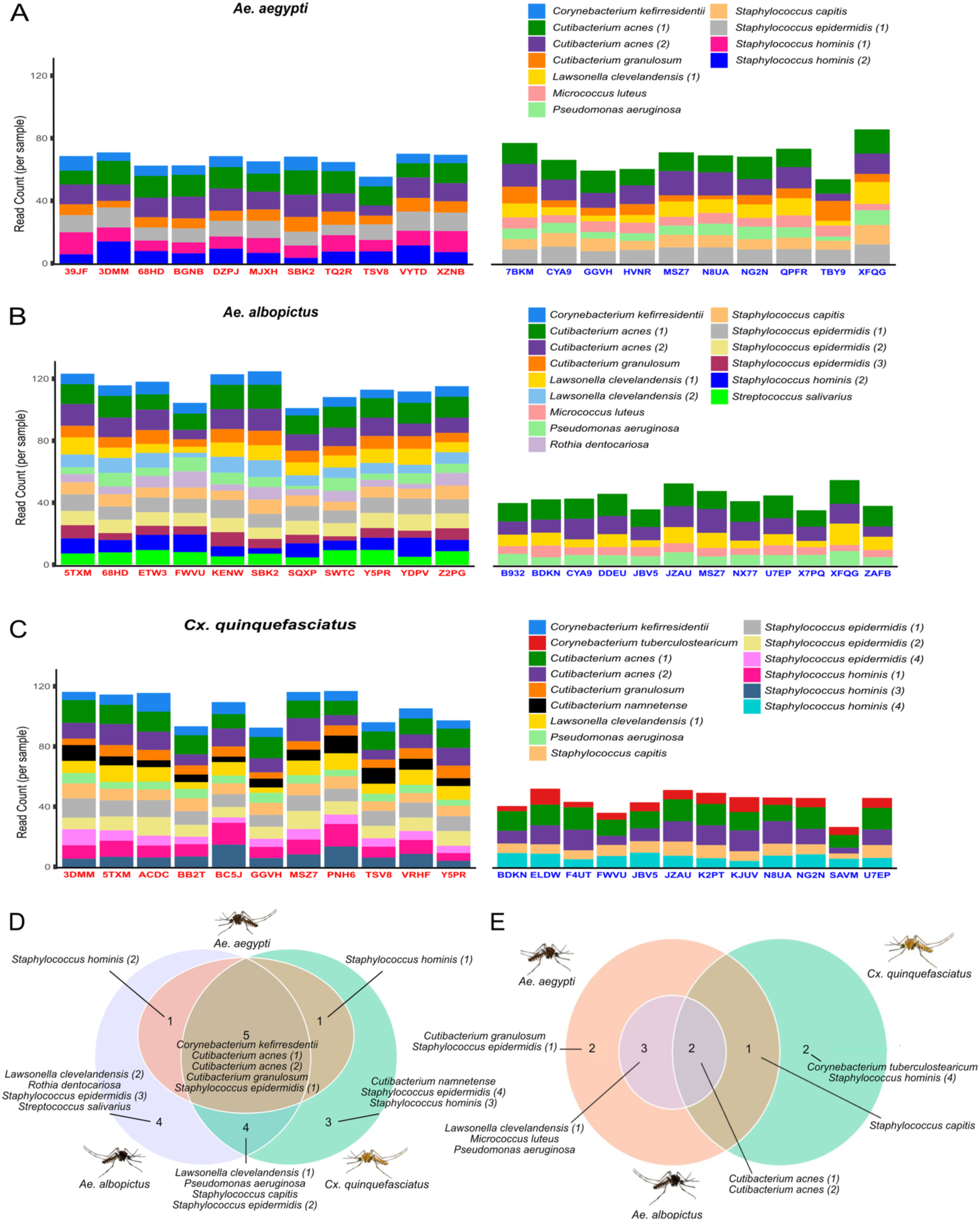
Core microbial communities reveal distinct and common taxa associated with species-specific mosquito attraction. **(A-C)** Stacked bar chart depicting the core skin commensals present for all high (left) or low (right) attraction participants for **(A)** *Ae. aegypti,* **(B)** *Ae. albopictus,* or **(C)** *Cx. quinquefasciatus.* Bacteria are labeled by ASV. **(D)** Venn diagram of core microbes for all high attraction participants across all mosquito species. **(E)** Venn diagram of core microbes for all low attraction participants across all mosquito species.

We did not detect any distinguishing core microbes in *Ae. aegypti* high attraction participants from other species tested, while *Ae. albopictus* high attraction participants had four distinct bacterial species and *Cx. quinquefasciatus* high attraction participants had three distinct bacterial species. There were only five bacteria that were associated with high attraction across all three mosquito species: *Corynebacterium kefirresdentii, C. acnes* (*1*)*, C. acnes* (*2*)*, C. granulosum,* and *S. epidermidis* (*1*) (Fig. 6D). No unique core microbes were detected in low attraction *Ae. albopictus* participants, but two distinct bacteria were detected in each of the *Ae. aegypti* and *Cx. quinquefasciatus* low attraction participants. Only two ASVs of *C. acnes* comprised the shared core microbiome for low attraction participants across all vector species tested (Fig. 6E). Across *Ae. aegypti*, *Ae. albopictus*, and *Cx. quinquefasciatus* mosquitoes, we detected three core bacteria in common for high attraction participants that were not found across the low attraction core microbiomes. These bacterial taxa, *Corynebacterium kefirresidentii*, *Cutibacterium granulosum*, and *Staphylococcus epidermidis,* may broadly signal to mosquitoes the presence of an attractive host.

## DISCUSSION

To determine species-specific odor and bacterial signatures that drive mosquito attraction to humans, we tested the attraction of 119 participants to three mosquito vector species: *Ae. aegypti, Ae. albopictus,* and *Cx. quinquefasciatus*. Our findings indicated sex-specific differences in *Ae. aegypti* mosquito attraction, but not in *Ae. albopictus* or *Cx. quinquefasciatus* mosquitoes. When we analyzed the odor profiles and microbiomes between male and female participants, we found many differences that may contribute to *Ae. aegypti* attraction rates. There was high diversity in individual skin microbiomes of the participants in our study. Strikingly, the core microbiomes of both male and female participants consisted of only one species of bacteria.

Male participants exhibited a significant increase in bacteria associated with the production of fatty acid volatiles, including the *Corynebacterium* genus, that may enhance the attraction of *Ae. aegypti* (53, 54). Notably, we found female participants showed higher abundances of *Pseudomonas aeruginosa,* which is consistent with previous work depicting a reduced attraction in both *An. gambiae* and *Ae. aegypti* mosquitoes (43, 44). Although we have identified many differences between male and female odor plumes and skin bacteria, we do not know which of these features make male participants more attractive than female participants to *Ae. aegypti* mosquitoes. Given this complexity, we sought to identify the cues that correlate with attraction for each of the three vector species.

We found that different odor plumes contributed to each mosquito species’ attraction rates to humans. *Ae. aegypti* and *Cx. quinquefasciatus* attraction was associated with the absence of repulsive odors in the odor plume. For *Ae. aegypti,* only one odor was more abundant in high attraction participants than low attraction participants, and there were no odors detected in these high attraction participants that distinguished them from the other species tested. Yet, eight compounds are species-specific for *Ae. aegypti* low attraction participants. These odors are alpha-fenchene, mesitylene, p-cymene, sabinene hydrate, (Z)-4-decen-1-ol, unknown (RI 1301), alpha-barbatene, and isobornyl butanoate. This lack of high attraction species-specific odors could also reflect the importance of odors interacting in a plume rather than individually. This species is almost exclusively anthropophilic, so we expect the olfactory receptors of these mosquitoes to be finely tuned for specific human cues (6, 15, 18, 56). Our evidence suggests the lack of repellent odors may cause *Ae. aegypti* mosquitoes to target a human host.

*Cx. quinquefasciatus* followed a similar pattern to *Ae. aegypti* and did not have any species-specific odors for high attraction but had five for low attraction. These were farnesane, isoborneol, 1,4-dichlorobenzene, benzaldehyde, and an unknown odor (RI 811). Behaviorally, these mosquitoes are more opportunistic and will blood feed on birds, pigs, and reptiles as well as humans (56–58). The association of high attraction participants with the absence of repulsive odors in *Cx. quinquefasciatus* suggests that these odors, some of which have been linked with repellency, allow this species to distinguish hosts that contain a broad set of potential attractants with a narrower set of repellent compounds.

Conversely, we found that *Ae. albopictus* attraction rates were associated with the presence of attractive odors. Along with strong preferences for cyclic alcohols and terpenoids, we found that *Ae. albopictus* had thirteen species-specific odors in high attraction participants that were not found in the other three species tested. Several of these odors are derived from plant-like environments and may reflect the generalist feeding behavior and ecological flexibility of *Ae. albopictus* (59, 60). Though it has been commonly viewed as an invasive species, *Ae. albopictus* have been shown to be opportunistic feeders (61–64). The cues that *Ae. albopictus* use for host-seeking are more closely related to ancestral, zoophilic *Ae. aegypti* rather than the human-specialized *Ae. aegypti* that inhabit urbanized environments (65). Therefore, it is likely to be evolving its own distinct path to human host detection, and our results support that *Ae. albopictus’* attraction is guided by a different set of cues than its sister species *Ae. aegypti* (66–68).

Both abundance and community composition of the human skin microbiome contribute to species-specific mosquito attraction via odor production (69–71). Biotransformation of the products of sweat and sebum produce VOCs, but community interactions also play a role in the resulting odor plume (29, 35, 49, 50, 72–74). For *Ae. aegypti* and *Ae. albopictus,* attraction may be driven by both the presence and absence of attractive and repellent bacterial taxa. Conversely, attraction rates of *Cx. quinquefasciatus* mosquitoes may be driven by the absence of repellent bacterial taxa. Unsurprisingly, *Pseudomonas spp.* was detected not only in higher abundance in *Ae. aegypti* low attraction participants, but also in the low attraction core microbiome. We have identified several species of *Pseudomonas* that are associated with low attraction for *Ae. aegypti*, including *Pseudomonas A stutzeri*, *Pseudomonas B oryzihabitans*, and *Pseudomonas aeruginosa.* This may indicate that *Pseudomonas spp.* is responsible for the production of repellent odors. In studies with *An. gambiae*, *Pseudomonas aeruginosa* has been associated with individuals that are poorly attractive (36). For both *An. gambiae* and *Ae. aegypti*, *in vitro* microbial communities with higher ratios of *Pseudomonas aeruginosa* were less attractive (36, 44). Additionally, we hypothesize that *Pseudomonas spp.* exhibit antimicrobial activity within the skin microbiome community, which could explain both its higher abundance and contribution to the odor plume (75–77). Microbes contributing to repulsive odors are distinct for *Cx. quinquefasciatus,* as only *A. oris* was more abundant in low attraction. This species has been used as antimicrobial larvicide in vector control strategies, so it may share similar interspecific bacterial interactions amongst the skin microbiome (78–80). The absence of distinct repellent odors is associated with increased attraction in both *Ae. aegypti* and *Cx. quinquefasciatus* mosquitoes, and the underlying skin microbes are also different. For example, the *Cx. quinquefasciatus* low attraction core microbiome does not have *Pseudomonas spp.,* reflecting the difference between *Ae. aegypti* and *Cx. quinquefasciatus* in the cues they likely detect when human host-seeking.

For *Ae. albopictus,* we hypothesize that the human skin bacteria are driving attraction by producing more attractive odors. We found that there were twice as many significantly abundant microbial species for high attraction participants than low attraction participants. Of interest was the *Anaerococcus spp.,* including *A. octavius,* which is linked to the production of volatile fatty acids through the breakdown of sweat components (81). It is likely that biotransformations such as these lead to attractive odor plumes that *Ae. albopictus* mosquitoes can detect.

How humans are generally perceived by mosquitoes is becoming clearer, but what determines whether an individual is targeted by a mosquito has been difficult to assess (6, 7, 11, 18, 39, 82–84). We have found that species-specific odor and bacteria signatures associated with individual humans are correlated with attraction rates by *Ae. aegypti, Ae. albopictus,* and *Cx. quinquefasciatus* mosquitoes. Thus, we have observed that different mosquito species are attracted to different human cues. Of the 3,500 mosquito species, only a few have developed anthropophily, and the evolutionary paths to this are likely to be distinct (85). Our data set provides perspective on how attraction to humans in three vector species can be driven by both distinct and common cues and is consistent with each species independently evolving their host preference. Unraveling this complexity will aid ongoing efforts to generate microbial-based mosquito repellents that produce repulsive compounds or reduce attractive cues (86). The bacterial and odor signatures provided may also be useful to interrupt mosquito interactions with humans.

## MATERIALS AND METHODS

### Recruitment

All research in this study was reviewed and approved by the Florida International University Institutional Review Board under protocol #IRB-16-0386-AM02. To characterize the signatures of human odor that drive mosquito attraction, we recruited, obtained informed consent, and successfully assayed 119 participants between the ages of 18-59. Participants self-reported their race and ethnicity, with White Hispanic/Latinx participants making up 42%, 21% identified as White non-Hispanic, 16% as Black, 15% as Multiracial, and 6% as Asian. The sex of participants was also self-reported, with 51% male participants and 49% female participants.

### Assessing the attractiveness of individual humans

Mosquitoes were placed in a custom-made uniport olfactometer to analyze their behavioral attraction (8, 46). The uniport olfactometer consists of a large plexiglass tube (75 cm long and 13 cm wide) connected to a small cylindrical cage (13 cm long and 5 cm in diameter) that contains the mosquitoes prior to the experiment. At the far end of the plexiglass tube connected to the stimulus chamber, the left arm of the participant is inserted in an enclosed space with dimensions of 25 cm by 20 cm by 13 cm. In the stimulus chamber, carbon-filtered, humidified air and CO_2_ can combine with odorants to attract mosquitoes that have been released from a trap. Acrylic flowmeter Model VFA-4-SSV (Dwyer Instruments Inc., IN, USA) set to 3 SCFH was used to measure the CO_2_ release rate in the stimulus chamber. The final concentration of CO_2_ in the assay was maintained at 2500-2700 ppm by a carbon dioxide monitor (Catalog#CO2-100, Amprobe). At the same time, the airflow rate was set at 21 standard cubic feet per hour by an air flowmeter (King Instruments CA, USA). The sealed design of the uniport, air filtration, and the positive pressure caused by air circulation in the apparatus will isolate the assay from all possible environmental scents.

Prior to experimentation, participants were instructed to refrain from showering the night before their visit and were heavily discouraged from wearing skin products such as deodorants, antiperspirants, perfume, or cologne. Uniport olfactometry was performed during volunteer visits for both *Ae. aegypti* and *Ae. albopictus*, in triplicate for each species, to determine attraction. To quantify *Cx. quinquefasciatus* attraction, participants wore nylon stockings over their arms for 12-16 hours to collect odor. The stockings were worn following their shower the evening before the study and were collected at the beginning of their visit and stored at −20°C until use in the olfactometer.

Each olfactometer experiment was conducted for eight minutes and contained 30 females of each species. Each participant underwent six trials of olfactometer assays, three with *Ae. aegypti* and three with *Ae. albopictus*. *Cx. quinquefasciatus* are night-biting mosquitoes, and trials were performed under moonlight (lux 0.05-0.1) conditions using nylon sleeves in the olfactometer rather than a live human. Before each human experiment, a blank (no human or nylon sleeve) trial is conducted with each species. During the investigation, the mosquito species is randomized. All mosquitoes were subjected to behavioral experiments only once before being sacrificed.

Mosquito attraction rates were calculated as done previously (47). All assays were initially set up with 30 mosquitoes in the WHO tube release chamber. Mosquitoes that left the release chamber and entered the attraction trap were scored as attracted. Mosquitoes that did not leave the release chamber were scored as unactivated and included in the mosquito total. Mosquitoes that were dead were removed from the mosquito total. The attraction rate was calculated by dividing the number of attracted mosquitoes by the total number of live mosquitoes.

### Profiling body odor collection

To identify volatile organic compounds (VOCs) emanating from the epidermis, we collected volatiles from the headspace of each participant’s left arm. The left arm was placed in a nylon bag (Toppits, Cofresco Frishhalteprodukte GmbH & Co., Minden, Germany), and their volatiles were collected for 90 minutes on Tenax adsorbent fibers by circulating ultra zero grade air (Airgas) at a rate of 400 mL min^-1^ per minute into the top of the bag and simultaneously applying a vacuum (GAST, Model DOA-P704-AA, Michigan, USA) to pull air at a 200 mL min^-1^ rate to the back of the stainless steel thermal desorption tubes filled with 200 mg of Tenax TA.

Using a GC-MS equipped with a thermodesorption unit, we characterized human volatile profiles. VOCs were desorbed from the Tenax tubes for 10 minutes at 250°C and captured in a sorbent trap chilled with liquid nitrogen at −110°C. Compounds were desorbed from this trap during secondary desorption at 40°C s^−1^ and desorbed at 280°C for 10 min, and later transferred in split mode to a non-polar gas-chromatography (GC) column (RXI-5ms 30 m×0.25 mm×1.00 μm). The chromatographic procedure was conducted at a carrier gas flow rate of 1 mL min^-1^. The temperature of the GC oven was programmed to rise from 40°C (5 min hold time) to 280°C (8 min hold time) at a rate of 5°C min^-1^. The temperature of the MS transfer line was set to 280°C. The electron beam energy was set to 70 eV, and the ion source temperature was set to 250°C. The mass spectrometer scanned m/z 35–400 at a rate of 4.7 scans s^−1^. Helium gas was used for desorption and chromatographic analyses. GC–MS data were processed using the MetAlign–MSClust software pipeline (87). In brief, MetAlign corrects the baseline and eliminates the noise of each GC–MS output file. Subsequently, it aligns the individual mass peaks in all chromatograms (88). MSClust then clusters the aligned mass peaks so that mass spectra of putative compounds are reconstructed (89).

### Sampling the human skin microbiome

Following behavioral analysis, the participant rested their arm atop a table covered with a sterile surgical drape. A total of four samples were collected from the underside of the individual’s arm, each comprising a 5 x 5 cm area. The sampling sites are located on the distal volar forearm (beginning just above the wrist crease), mid-volar forearm, proximal volar forearm, and antecubital fossa.

For each sample site, two sterile double swabs (BD BB CultureSwab EZ II) were used to collect samples. Each swab was moistened with a sterile aliquot of SCF-1 buffer (50 mM Tris buffer [pH 7.6], 1 mM EDTA [pH 8.0], and 0.5% Tween-20) and vigorously swabbed against the skin for two minutes, rolling swabs to collect along the entire flocked surface. Swabs from each sample site were clipped with sterilized cutters into a 25 ml conical tube containing 1 ml of DNA/RNA Shield (Zymo Research, California, USA) and mixed. Blank samples were collected for each participant by waving swabs moistened with SCF-1 buffer in the air for two minutes, then collected and processed identically in DNA/RNA Shield.

For extraction, samples were first vortexed for 1 minute to displace cells from the swab to the DNA/RNA Shield. The suspension was then transferred to a BashingBead Lysis Tube containing 0.1 & 0.5 mm glass beads (Zymo Research, California, USA). Sample suspensions were then lysed in the ZR BashingBead Lysis tube (0.1 & 0.5 mm beads) in DNA/RNA Shield using the Omni Bead Rupto-24 Elite (Omni International, Kennesaw, GA) (velocity of 6 m/s for 1 minute, 5 minutes rest, cycle is repeated three times for a total of 3 minutes of bead beating). DNA was extracted from lysed samples using the ZymoBIOMICS DNA Microprep Kit (Zymo Research). DNA samples were amplified with indexed 27F (5’GCATC/barcode/AGRGTTYGATYMTGGCTCAG3’) and 1492R (5’GCATC/barcode/RGYTACCTTGTTACGACTT3’) primers, with 16 base pair barcodes used in unique combinations to identify libraries. Libraries were constructed in a 25 μl reaction volume with 2.5 μM of each primer and 12.5 μ KAPA HiFi HotStart ReadyMix PCR Kit (KAPA Biosystems). Libraries were then amplified using the following program: a 95°C initial denaturation for three minutes, followed by 28 cycles of amplification with 95°C denaturation for 30 seconds, 57°C annealing for 30 seconds, and 30-second extension at 72°C, with a final extension step at 72°C for 60 seconds. Libraries were purified on AMPure PB beads (Pacific Biosciences), quantified via Qubit dsDNA HS (Thermo Fisher), and normalized prior to pooling. SMRTbell libraries were constructed from amplicon pools and sequenced for a 15-hour movie duration on the Sequel II Platform (Pacific Biosciences). Participant human skin microbiomes were characterized with full-length (V1-V9) 16S amplicon sequencing using PacBio HiFi sequencing. Previous mosquito/host microbiome investigations used short fragment Illumina 16S sequencing, however, our method captures all the hypervariable areas of 16S, vastly improving the assay’s taxonomic accuracy. Demultiplexed libraries were denoised to amplicon sequence variants (ASVs) using DADA2 (90) and analyzed using QIIME 2 (91), with classification performed using VSEARCH (92) in combination with the Silva 138 99% OTUs full-length sequences database. The ASV method begins by identifying which precise sequences were read and how often each precise sequence was read. These data are then combined with an error model for the sequencing run, allowing the comparison of similar readings to determine the probability that a particular read at a particular frequency is not due to sequencing error and subsequently filtered for PCR chimeras (93). Essentially, this generates a p-value for each exact sequence, where the null-hypothesis is equivalent to that exact sequence being the result of sequencing error (90).This resulted in the species-level identification of nearly all major taxa present in the samples. Occasionally, this method can identify fungal species via off-target sequencing or non-specific amplification of mitochondrial DNA, and so any detected fungal species were removed from the analysis (94–96).

### Statistical analysis

#### Identifying high and low attraction participants

High and low attraction participants for *Ae. aegypti, Ae. albopictus*, and *Cx. quinquefasciatus* were identified through a percentile analysis to isolate participants in the top 10% (90^th^ percentile) and bottom 10% (10^th^ percentile) of ranked attraction for each vector species. This gave us 13 high attraction participants and 12 low attraction participants for *Ae. aegypti,* 12 high and low attraction participants for *Ae. albopictus*, and 12 high and low attraction participants for *Cx. quinquefasciatus.* We then compared both high and low attraction groups for each vector species with a Wilcoxon rank-sum test, and both groups significantly differed from each other (*P* < 0.0001).

#### Identification of microbial components

The microbiome data were analyzed using the Quantitative Insights into Microbial Ecology 2 (QIIME2) (version 2019.1) program. The feature table, phylogeny data, and metadata table were then imported for analysis into MicrobiomeAnalyst (version 2.0) (97). Participants 3AXB, RPYF, SNTZ, AKFQ, TAFA, and WRQX did not have microbiome sequencing data, so these participants were excluded from all microbiome analyses. Based on the mean abundance of operational taxonomical units (OTUs), samples were filtered for low prevalence (removed if 10% of counts were less than 4 across all samples) and 10% variability using a rank-based inter-quantile range assessment. The data was normalized using cumulative sum scaling (CSS). Stacked column graphs were constructed with this software using the 16S reads as relative abundance. The participants of each group made up each column (the x-axis), and total bacterial counts made the y-axis. Stacks in the columns are constructed from the read counts for each OTU in the microbiome for the group. A Wilcoxon rank-sum heat tree was created to visualize differences between high attraction and low attraction groups for each vector species.

#### HA/LA core microbiome groups for mosquito species

The level of attraction to each of the three mosquito species (*Ae. aegypti, Ae. albopictus,* or *Cx. quinquefasciatus*) was matched with each participant’s microbiome sequencing data so that groups that are high attraction or low attraction to each species were formed. Using the *phyloseq* R package, physeq objects were created for each mosquito species by combining the feature table, phylogeny data, and metadata. Each of these physeq objects were filtered for a minimum of four read counts in 10% of the data and for a rank-sum based inter-quantile variance of 10%.

Data were then scaled using cumulative sum scaling (CSS). Zeroes from the data were dropped, and the groups in the physeq object were separated by the metadata to form distinct, filtered, and scaled high- and low-attraction groups for each vector species. Next, each group was assessed for bacteria that are present across every participant in the group (e.g., bacteria that are present in all participants who are highly attractive to *Ae. aegypti*). These bacteria were classified at the amplicon sequence variant (ASV) level, and the scaled counts of reads were recorded. Finally, stacked column graphs were constructed using the *ggplot2* package, where the volunteers in each group made up each column (the x-axis), and the total microbial counts made the y-axis. Stacks in the columns are constructed from the read counts for each ASV in the core microbiome for the group. All data analysis was performed using *R/RStudio*.

#### Male/Female core microbiome for mosquito species

Participants were matched with their reported biological sex and their microbiome sequencing data so that male and female groups were formed. Using the *phyloseq* R package, physeq objects were created for each mosquito species by combining the feature table, phylogeny data, and metadata. Each of these physeq objects were filtered for a minimum of four read counts in 10% of the data and for a rank-sum based inter-quantile variance of 10%. Data was then scaled using cumulative sum scaling (CSS). Zeroes from the data were dropped, and the groups in the physeq object were separated by the metadata to form distinct, filtered, and scaled male and female groups. Next, each group was assessed for bacteria that were present across every participant in the group (e.g., bacteria that were present in every male in the study). These bacteria were classified at the amplicon sequence variant (ASV) level, and the scaled counts of reads were recorded. Finally, stacked column graphs were constructed using the *ggplot2* package, where the volunteers in each group made up each column (the x-axis), and the total bacterial counts made the y-axis. Stacks in the columns are constructed from the read counts for each ASV in the core microbiome for the group. All data analysis was performed using *R/RStudio*.

## DATA AVAILABILITY

The DNA sequencing data files are available for download on NCBI Sequence Read Archive (PRJNA1415315). All underlying data used to generate figures in this manuscript as well as R scripts for statistical analyses and visualization are available via dryad data depository (DOI: 10.5061/dryad.vdncjsz97).

## FUNDING

Defense Advanced Research Projects Agency (DARPA) Biological Technologies Office (BTO) (HR0011-20-C-0073) (JSC, AJB, DC, ALCS, NOV, MD).

Centers for Disease Control and Prevention (CDC), Southeastern Center of Excellence in Vector-borne Disease (U01CK000662) (KMM).

Florida International University Graduate Teaching (MAM).

## ACKNOWLEDGEMENTS

We would like to thank the DeGennaro lab for their support and feedback on the manuscript. We thank Kristian Lopez for his technical contributions to the manuscript. We also appreciate the suggestions Takeshi Morita and Sheng-Hao Lin provided for the manuscript.

## AUTHOR CONTRIBUTIONS

Conceptualization: NOV and MD. Methodology: JSC, KMM, DLB, AJB, DC, ALCS, NOV, and MD. Software: KMM, AJB, and MAM. Formal Analysis: KMM, JSC, DLB, AJB, and MAM. Investigation: JSC, KMM, DLB, AJB, DC, and ALCS. Validation: JSC, KMM, ALCS, and MD. Resources: NOV and MD. Data Curation: JSC, KMM, ALCS, and MD. Writing – original draft: KMM, JSC, and MD. Writing – review and editing: KMM, DLB, AJB, DC, MAM, ALCS, NOV, and MD. Visualization: JSC, KMM, MAM, and MD. Project Administration: NOV and MD. Funding Acquisition: NOV and MD.

## SUPPLEMENTAL MATERIAL

**Figure S1.**
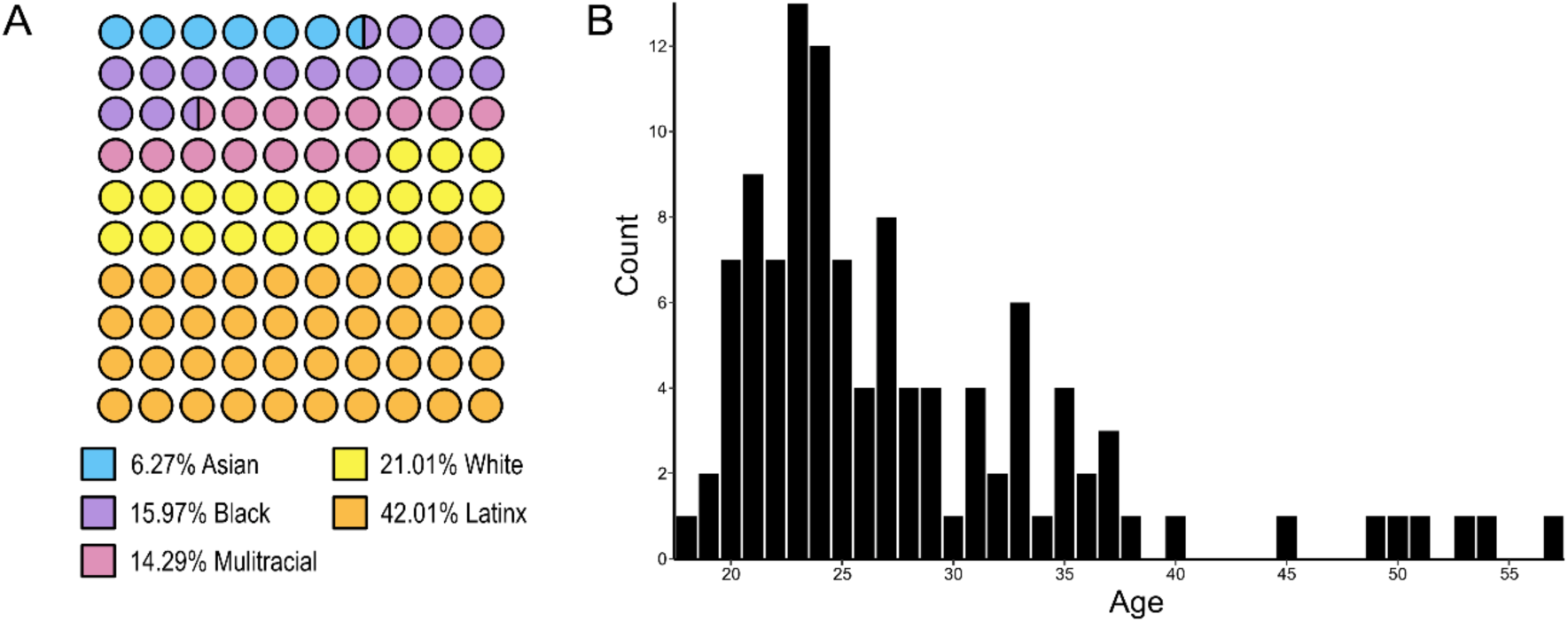
**(A)** Distribution of self-reported race for the 119 participants in this study. **(B)** Histogram showing age distributions of the participants in the study.

**Figure S2.**
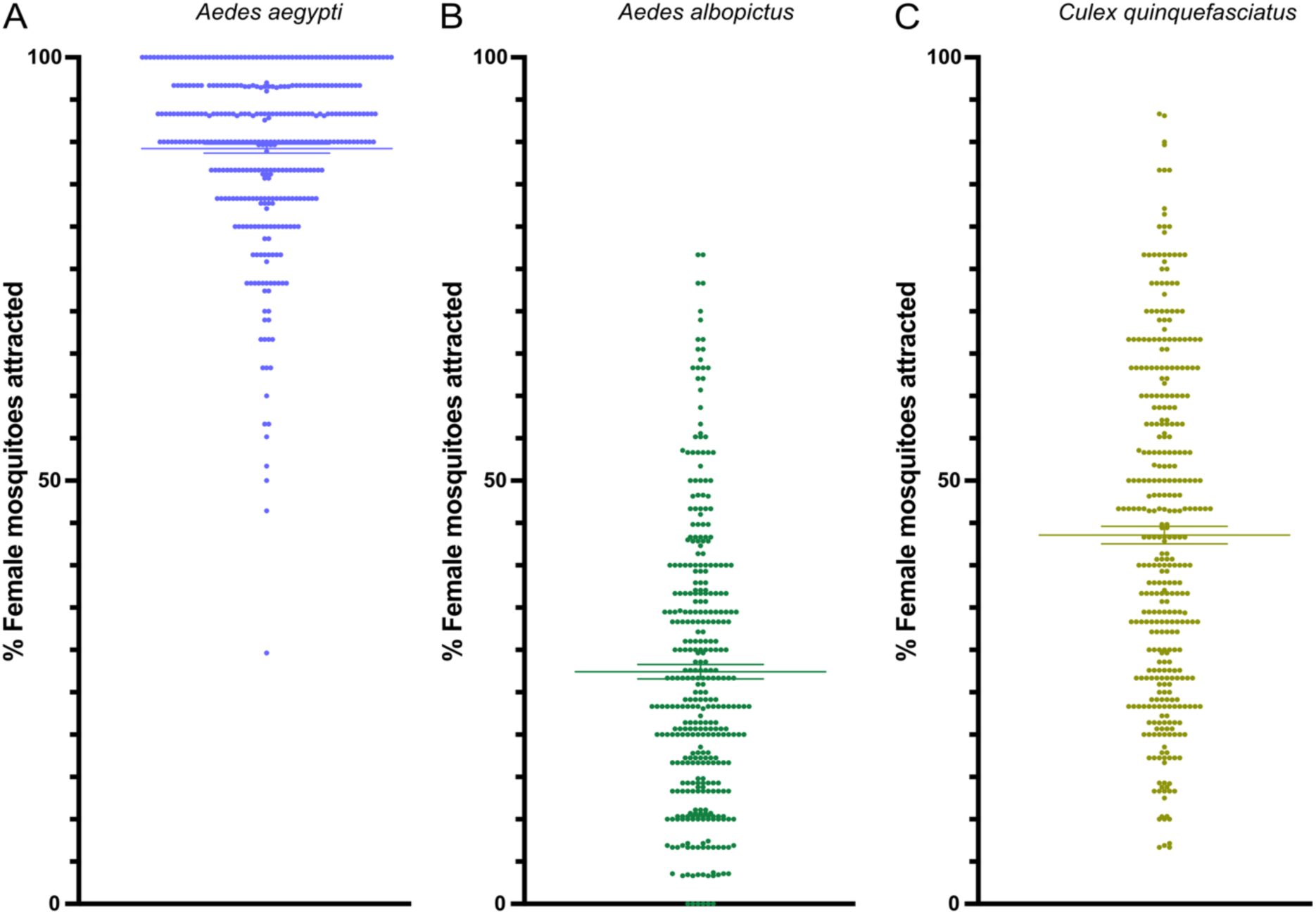
Overall attraction rates of **(A)** *Aedes aegypti,* **(B)** *Aedes albopictus,* and **(C)** *Culex quinquefasciatus* female mosquitoes in uniport olfactometer experiments. The y-axis represents the percentage of female mosquitoes attracted, while data points for each group show individual values, with horizontal lines indicating the standard error of the mean.

**Figure S3.**
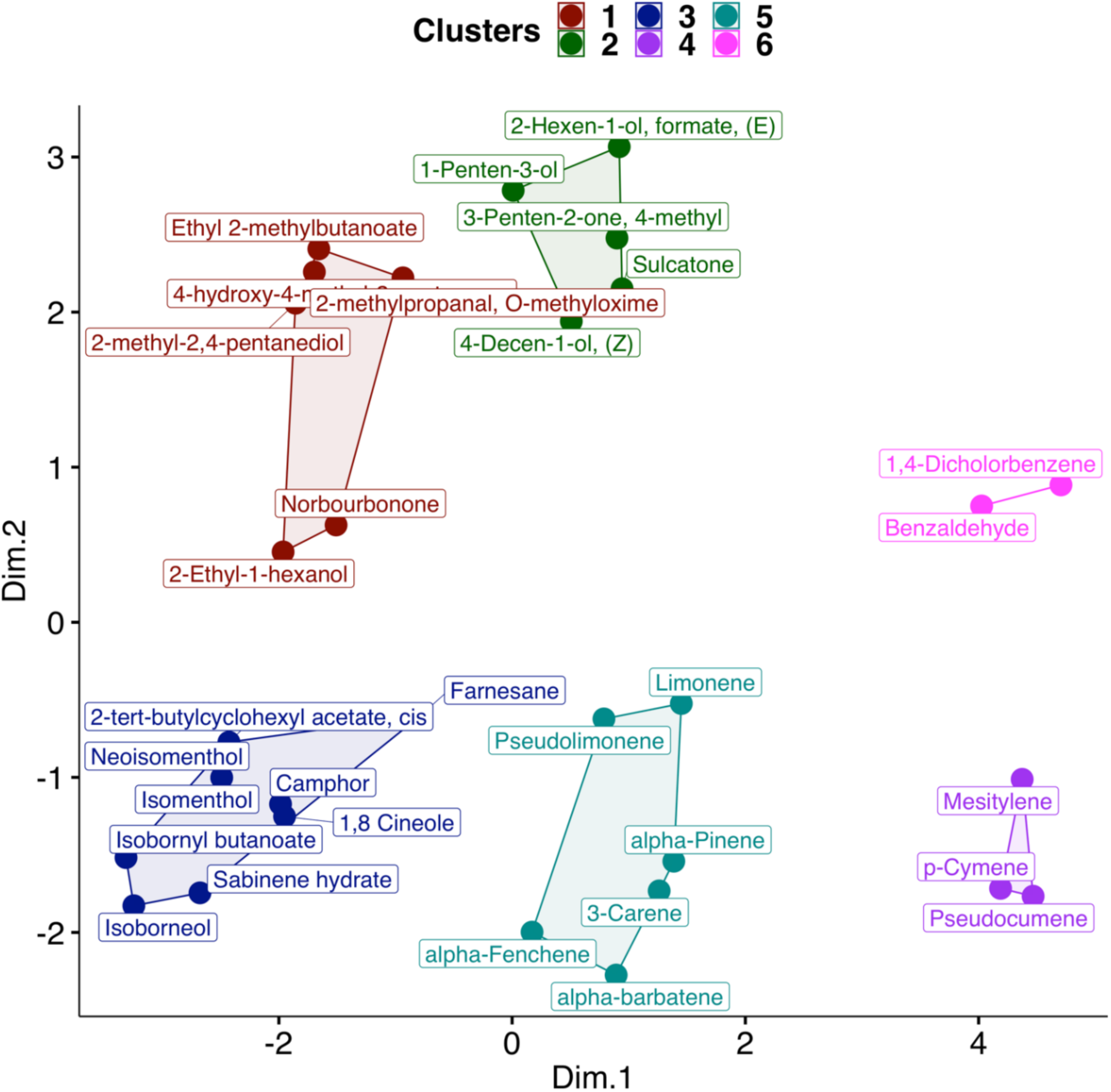
K-means clustering of chemicals based on their atomic structure, plotted along two principal dimensions from multidimensional scaling (MDS). Chemicals are color-coded by cluster: Cluster 1 (red) includes oxygenated and branched hydrocarbons like Ethyl 2-methylbutanoate; Cluster 2 (green) contains unsaturated alcohols and ketones such as 2-Hexen-1-ol, formate (E); Cluster 3 (blue) features cyclic alcohols and terpenoids like Isoborneol; Cluster 4 (purple) includes methyl-substituted aromatics such as Mesitylene; Cluster 5 (cyan) highlights monoterpenes like Limonene and alpha-Pinene; and Cluster 6 (pink) groups aromatic chemicals like Benzaldehyde. Cluster separation reflects structural similarities, with proximity indicating chemical relationships within the principal dimensions.

**Figure S4.**
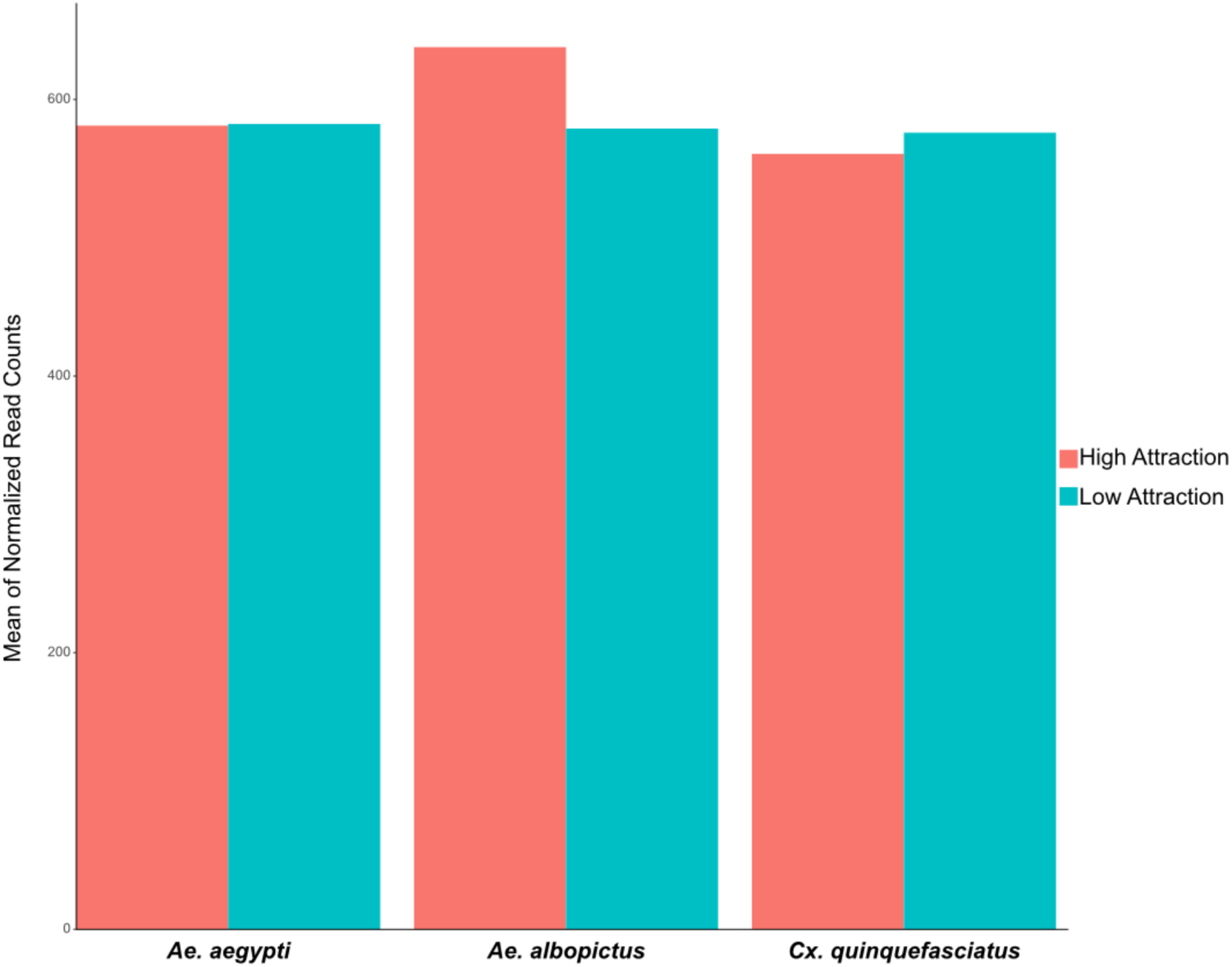
Mean normalized read counts of human skin microbiome for high and low attraction groups in each mosquito species. *Ae. aegypti* high attraction had an average of 581.115 and low attraction had an average of 582.296. *Ae. albopictus* had a high attraction average of 637.834 and a low attraction average of 578.947. *Cx. quinquefasciatus* high attraction had an average of 560.657 and low attraction had an average of 575.998.

**Figure S5.**
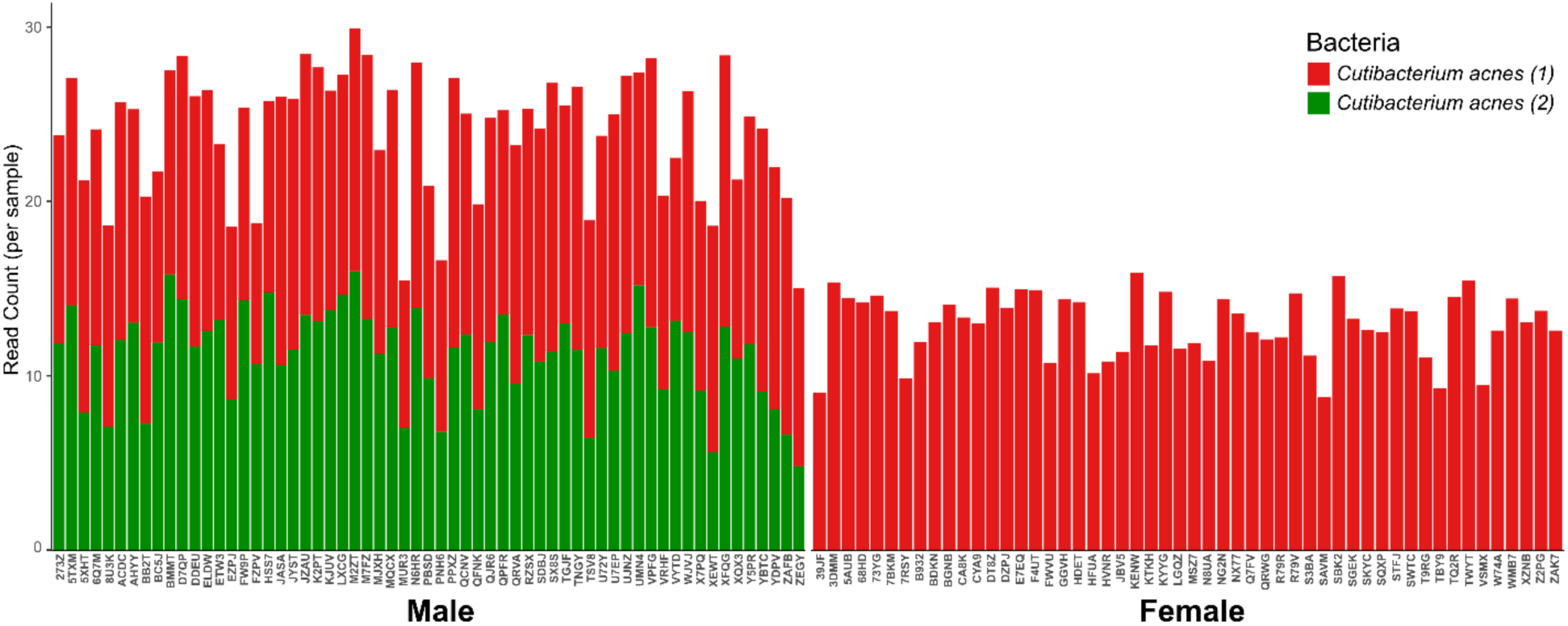
Core microbiome of all bacteria detected for each self-reported male (left) and female (right) individual. Two amplicon sequence variants (ASVs) of *C. acnes* comprise the male core microbiome, while one ASV of *C. acnes* comprises the female core microbiome. All bacteria were normalized and scaled using cumulative sum scaling.

**Table S1.** Volatile organic compounds were identified via GC-MS and evaluated for greater prevalence in high attraction or low attractions groups for each mosquito species. Text in red denotes that the odor was more abundant in the high attraction group for that mosquito species. Text in blue denotes that the odor was more abundant in the low attraction group for that mosquito species.

| Odor | *Ae. aegypti* | *Ae. albopictus* | *Cx. quinquefasciatus* |
| --- | --- | --- | --- |
| Unknown (RI 811) | High Attraction ▲ | High Attraction ▲ | Low Attraction ▲ |
| Ethyl 2-methylbutanoate | High Attraction ▲ | High Attraction ▲ | High Attraction ▲ |
| (E)-2-Hexen-1-ol, formate | High Attraction ▲ | High Attraction ▲ | High Attraction ▲ |
| alpha-Pinene | High Attraction ▲ | High Attraction ▲ | High Attraction ▲ |
| Benzaldehyde | High Attraction ▲ | High Attraction ▲ | Low Attraction ▲ |
| Pseudolimonene | High Attraction ▲ | Low Attraction ▲ | High Attraction ▲ |
| 2-Ethyl-1-hexanol | High Attraction ▲ | Low Attraction ▲ | High Attraction ▲ |
| 1,4-Dichlorobenzene | High Attraction ▲ | High Attraction ▲ | Low Attraction ▲ |
| Isoborneol | High Attraction ▲ | High Attraction ▲ | Low Attraction ▲ |
| Farnesane | High Attraction ▲ | High Attraction ▲ | Low Attraction ▲ |
| 1-Penten-3-ol | Low Attraction ▲ | Low Attraction ▲ | Low Attraction ▲ |
| 4-methyl-3-Penten-2-one | Low Attraction ▲ | High Attraction ▲ | Low Attraction ▲ |
| Unknown (RI 834) | Low Attraction ▲ | High Attraction ▲ | Low Attraction ▲ |
| 4-Hydroxy-4-methyl-2-pentanone | Low Attraction ▲ | Low Attraction ▲ | Low Attraction ▲ |
| 2-Methyl-2,4-pentanediol | Low Attraction ▲ | High Attraction ▲ | Low Attraction ▲ |
| 2-Methylpropanal, O-methyloxime | Low Attraction ▲ | Low Attraction ▲ | Low Attraction ▲ |
| alpha-Fenchene | Low Attraction ▲ | High Attraction ▲ | High Attraction ▲ |
| Mesitylene | Low Attraction ▲ | High Attraction ▲ | High Attraction ▲ |
| Pseudocumene | Low Attraction ▲ | High Attraction ▲ | Low Attraction ▲ |
| Sulcatone | Low Attraction ▲ | High Attraction ▲ | Low Attraction ▲ |
| 3-Carene | Low Attraction ▲ | Low Attraction ▲ | Low Attraction ▲ |
| p-Cymene | Low Attraction ▲ | High Attraction ▲ | High Attraction ▲ |
| Limonene | Low Attraction ▲ | High Attraction ▲ | Low Attraction ▲ |
| 1,8-Cineole | Low Attraction ▲ | High Attraction ▲ | Low Attraction ▲ |
| Unknown (RI 1073) | Low Attraction ▲ | High Attraction ▲ | Low Attraction ▲ |
| Sabinene hydrate | Low Attraction ▲ | High Attraction ▲ | High Attraction ▲ |
| Camphor | Low Attraction ▲ | High Attraction ▲ | Low Attraction ▲ |
| Isomenthol | Low Attraction ▲ | Low Attraction ▲ | Low Attraction ▲ |
| Neoisomenthol | Low Attraction ▲ | High Attraction ▲ | Low Attraction ▲ |
| (Z)-4-Decen-1-ol | Low Attraction ▲ | High Attraction ▲ | High Attraction ▲ |
| Unknown (RI 1301) | Low Attraction ▲ | High Attraction ▲ | High Attraction ▲ |
| cis-2-tert-Butylcyclohexyl acetate | Low Attraction ▲ | High Attraction ▲ | Low Attraction ▲ |
| Unknown (RI 1344) | Low Attraction ▲ | High Attraction ▲ | Low Attraction ▲ |
| alpha-Barbatene | Low Attraction ▲ | High Attraction ▲ | High Attraction ▲ |
| Isobornyl butanoate | Low Attraction ▲ | High Attraction ▲ | High Attraction ▲ |
| Norbourbonone | Low Attraction ▲ | High Attraction ▲ | Low Attraction ▲ |

**Data S1.** Difference table showing significant bacteria by abundance between males and females for Wilcoxon rank-sum heat tree. Significant features are highlighted in orange.

**Data S2.** Difference table showing significant bacteria by abundance between *Ae. aegypti* high and low attraction groups for Wilcoxon rank-sum heat tree. Significant features are highlighted in orange.

**Data S3.** Difference table showing significant bacteria by abundance between *Ae. albopictus* high and low attraction for Wilcoxon rank-sum heat tree. Significant features are highlighted in orange.

**Data S4.** Difference table showing significant bacteria by abundance between *Cx. quinquefasciatus* high and low attraction for Wilcoxon rank-sum heat tree. Significant features are highlighted in orange.

## REFERENCES

1. Gabiane G, Yen PS, Failloux AB. Aedes mosquitoes in the emerging threat of urban yellow fever transmission. Reviews in Medical Virology. 32 (4):e2333. (2022).

2. Hale GL. Flaviviruses and the traveler: Around the world and to your stage. A review of West Nile, Yellow Fever, Dengue, and Zika viruses for the practicing pathologist. 36 (6):e100188. (2023).

3. Gould CV, Staples JE, Guagliardo SAJ, et. al. West Nile Virus: A review. Journal of the American Medical Association. 334 (7):618–628. (2025).

4. Gorris ME, Bartlow AW, Pitts T, Manore CA. Projections of *Aedes* and *Culex* mosquitoes across North and South America in response to climate change. 17 e100317. (2024).

5. Dekker T, Cardé RT. Moment-to-moment flight manoeuvres of the female yellow fever mosquito ( *Aedes aegypti* L.) in response to plumes of carbon dioxide and human skin odour. Journal of Experimental Biology. 214 (20):3480–94. (2011).

6. DeGennaro M, McBride CS, Seeholzer L, Nakagawa T, Dennis EJ, Goldman C, et al. *orco* mutant mosquitoes lose strong preference for humans and are not repelled by volatile DEET. Nature. 498 (7455):487–91. (2013).

7. McMeniman CJ, Corfas RA, Matthews BJ, Ritchie SA, Vosshall LB. Multimodal Integration of carbon dioxide and other sensory cues drives mosquito attraction to humans. Cell. 156 (5):1060–71. (2014).

8. Raji JI, Melo N, Castillo JS, Gonzalez S, Saldana V, Stensmyr MC, et al. *Aedes aegypti* mosquitoes detect acidic volatiles found in human odor using the IR8a pathway. Current Biology. 29 (8):1253–1262.e7. (2019).

9. Zhan Y, Alonso San Alberto D, Rusch C, Riffell JA, Montell C. Elimination of vision-guided target attraction in Aedes aegypti using CRISPR. Current Biology. 31 (18):4180–4187.e6. (2021).

10. Alonso San Alberto D, Rusch C, Zhan Y, Straw AD, Montell C, Riffell JA. The olfactory gating of visual preferences to human skin and visible spectra in mosquitoes. Nature Communications. 4;13(1):555. (2022).

11. Laursen WJ, Budelli G, Tang R, Chang EC, Busby R, Shankar S, et al. Humidity sensors that alert mosquitoes to nearby hosts and egg-laying sites. Neuron. 111(6):874–887.e8. (2023).

12. Morita T, Lyn NG, Von Heynitz RK, Goldman OV, Sorrells TR, DeGennaro M, et al. Cross-modal sensory compensation increases mosquito attraction to humans. Science Advances. 11,eadn5758 (2025).

13. Mukwaya LG. The role of olfaction in host preference by *Aedes* (Stegomyia) *simpsoni* and *Ae. aegypti*. Physiological Entomology. 1(4):271–6. (1976).

14. Lindsay SW, Adiamah JH, Miller JE, Pleass RJ, Armstrong JRM. Variation in attractiveness of human subjects to malaria mosquitoes (Diptera: Culicidae) in the Gambia. Journal of Medical Entomology.30 (2):368–73. (1993).

15. Takken W, Verhulst NO. Host preferences of blood-feeding mosquitoes. Annual Review of Entomology. 58 (1):433–53. (2013).

16. Dormont L, Bessière JM, Cohuet A. Human skin volatiles: a review. Journal of Chemical Ecoologyl. 39 (5):569–78. (2013).

17. Mukabana WR, Takken W, Coe R, Knols BG. Host-specific cues cause differential attractiveness of Kenyan men to the African malaria vector *Anopheles gambiae*. Malaria Journal.1 (1):17. (2002).

18. McBride CS, Baier F, Omondi AB, Spitzer SA, Lutomiah J, Sang R, et al. Evolution of mosquito preference for humans linked to an odorant receptor. Nature.515 (7526):222–7. (2014).

19. McBride CS. Genes and odors underlying the recent evolution of mosquito preference for humans. Current Biology. 26 (1):R41–6. (2016).

20. Zhao Z, Zung JL, Hinze A, Kriete AL, Iqbal A, Younger MA, et al. Mosquito brains encode unique features of human odour to drive host seeking. Nature. 605 (7911):706–12. (2022).

21. Knols BGJ, De Jong R, Takken W. Differential attractiveness of isolated humans to mosquitoes in Tanzania. Transactions of the Royal Society of Tropical Medicine and Hygiene.89 (6):604–6. (1995).

22. Qiu YT, Smallegange RC, Van Loon JJA, Ter Braak CJF, Takken W. Interindividual variation in the attractiveness of human odours to the malaria mosquito *Anopheles gambiae s. s*. Medical Vet Entomology.20 (3):280–7. (2006).

23. Gallagher M, Wysocki CJ, Leyden JJ, Spielman AI, Sun X, Preti G. Analyses of volatile organic compounds from human skin. British Journal of Dermatology. 159 (4):780–91. (2008).

24. Amann A, Costello BDL, Miekisch W, Schubert J, Buszewski B, Pleil J, et al. The human volatilome: volatile organic compounds (VOCs) in exhaled breath, skin emanations, urine, feces and saliva. Journal of Breath Research. 8 (3):034001. (2014).

25. Drabińska N, Flynn C, Ratcliffe N, Belluomo I, Myridakis A, Gould O, et al. A literature survey of all volatiles from healthy human breath and bodily fluids: the human volatilome. Journal of Breath Research. 15 (3):034001. (2021).

26. Shelley WB. Axillary odor: experimental study of the role of bacteria, apocrine sweat, and deodorants. AMA Archives of Dermatology and Syphilology.68 (4):430. (1953).

27. Leyden JJ, McGinley KJ, Hölzle E, Labows JN, Kligman AM. The microbiology of the human axilla and its relationship to axillary odor. Journal of Investigative Dermatology. 77 (5):413–6. (1981).

28. Meijerink J, Braks MAH, Brack AA, Adam W, Dekker T, Posthumus MA, et al. Identification of olfactory stimulants for *anopheles gambiae* from human sweat samples. Journal of Chemical Ecology. 26 (6):1367–82. (1999).

29. James AG, Casey J, Hyliands D, Mycock G. Fatty acid metabolism by cutaneous bacteria and its role in axillary malodour. World Journal of Microbiology and Biotechnology. 20 (8):787– 93. (2004).

30. Scharschmidt TC, Fischbach MA. What lives on our skin: ecology, genomics and therapeutic opportunities of the skin microbiome. Drug Discovery Today: Disease Mechanisms. 10 (3–4):e83–9. (2013).

31. Zung JL, McBride CS. Sebaceous origins of human odor. Current Biology. 35 (8):R303– 13. (2025).

32. Verhulst NO, Beijleveld H, Knols BG, Takken W, Schraa G, Bouwmeester HJ, et al. Cultured skin microbiota attracts malaria mosquitoes. Malaria Journal.8 (1):302. (2009).

33. Byrd AL, Belkaid Y, Segre JA. The human skin microbiome. Natural Reviews Microbiology. 16 (3):143–55. (2018).

34. Cundell AM. Microbial ecology of the human skin. Microbial Ecology. 76 (1):113–20. (2018).

35. Swaney MH, Nelsen A, Sandstrom S, Kalan LR. Sweat and sebum preferences of the human skin microbiota. Microbiology Spectrum.11 (1):e04180–22. (2023).

36. Verhulst NO, Andriessen R, Groenhagen U, Bukovinszkiné Kiss G, Schulz S, Takken W, et al. Differential attraction of malaria mosquitoes to volatile blends produced by human skin bacteria. PLoS ONE.5 (12):e15829. (2010).

37. Verhulst NO, Beijleveld H, Qiu YT, Maliepaard C, Verduyn W, Haasnoot GW, et al. Relation between HLA genes, human skin volatiles and attractiveness of humans to malaria mosquitoes. Infection, Genetics and Evolution.18:87–93. (2013).

38. Showering A, Martinez J, Benavente ED, Gezan SA, Jones RT, Oke C, et al. Skin microbiome alters attractiveness to *Anopheles* mosquitoes. BMC Microbiology. 22 (1):98. (2022).

39. De Obaldia ME, Morita T, Dedmon LC, Boehmler DJ, Jiang CS, Zeledon EV, et al. Differential mosquito attraction to humans is associated with skin-derived carboxylic acid levels. Cell. 185 (22):4099–4116.e13. (2022).

40. Zhang H, Zhu Y, Liu Z, Peng Y, Peng W, Tong L, et al. A volatile from the skin microbiota of flavivirus-infected hosts promotes mosquito attractiveness. Cell. 185 (14):2510–2522.e16. (2022).

41. Giraldo, D. et al. Human scent guides mosquito thermotaxis and host selection under naturalistic conditions. Current Biology 33, 2367–2382.e7 (2023).

42. Braks MAH, Takken W. Incubated human sweat but not fresh sweat attracts the malaria mosquito *Anopheles gambiae sensu stricto*. Journal of Chemical Ecology.25 (3):663–72. (1999).

43. Verhulst NO, Qiu YT, Beijleveld H, Maliepaard C, Knights D, Schulz S, et al. Composition of human skin microbiota affects attractiveness to malaria mosquitoes. PLoS ONE. 6 (12):e28991. (2011).

44. Lucas-Barbosa D, Balvers C, Bellantuono AJ, Castillo JS, Costa-da-Silva AL, De Moraes CM, et al. Competition matters: using *in vitro* community models to study the impact of human skin bacteria on mosquito attraction. Frontiers in Ecology and Evolution. 11:1156311. (2023).

45. Coutinho-Abreu IV, Jamshidi O, Raban R, Atabakhsh K, Merriman JA, Akbari OS. Identification of human skin microbiome odorants that manipulate mosquito landing behavior. Scientific Reports. 14, 1631 (2024).

46. Castillo JS, Bellantuono AJ, DeGennaro M. Building a uniport olfactometer to assess mosquito responses to odors. Cold Spring Harbor Protocols. 2023 (10):pdb.prot108174. (2023).

47. Castillo JS, Bellantuono AJ, DeGennaro M. Quantifying mosquito attraction using a uniport olfactometer. Cold Spring Harbor Protocols. 2023 (10):pdb.prot108175. (2023).

48. Xie L, Yang W, Liu H, Liu T, Xie Y, Lin F, et al. Enhancing attraction of the vector mosquito *Aedes albopictus* by using a novel synthetic odorant blend. Parasites Vectors. 12 (1):382. (2019).

49. Grice EA, Kong HH, Conlan S, Deming CB, Davis J, Young AC, et al. Topographical and temporal diversity of the human skin microbiome. Science.324 (5931):1190–2. (2009).

50. Costello EK, Lauber CL, Hamady M, Fierer N, Gordon JI, Knight R. Bacterial community variation in human body habitats across space and time. Science. 326 (5960):1694–7. (2009).

51. Kim D, Crippen TL, Jordan HR, Tomberlin JK. Quorum sensing gene regulation in *Staphylococcus epidermidis* reduces the attraction of *Aedes aegypti* (L.) (Diptera: Culicidae). Frontiers in Microbiology. 14:1208241. (2023).

52. Campodónico VL, Llosa NJ, Grout M, Döring G, Maira-Litrán T, Pier GB. Evaluation of flagella and flagellin of *Pseudomonas aeruginosa* as vaccines. Infection and Immunity. 78 (2):746–55. (2010).

53. Schweizer E, Hofmann J. Microbial Type I Fatty Acid Synthases (FAS): Major Players in a Network of Cellular FAS Systems. Microbiology and Molecular Biology Reviews. 68 (3):501– 17. (2004).

54. Wang J, Murphy EJ, Nix JC, Jones DNM. *Aedes aegypti* Odorant Binding Protein 22 selectively binds fatty acids through a conformational change in its C-terminal tail. Scientific Reports. 10 (1):3300. (2020).

55. Harrington LC, Edman JD, Scott TW. Why do female *Aedes aegypti* (Diptera: Culicidae) feed preferentially and frequently on human blood? Journal of Medical Entomology. 38 (3):411– 22. (2001).

56. Bhattacharya S, Basu P. The Southern House Mosquito, *Culex quinquefasciatus*: profile of a smart vector. Journal of Entomology and Zoology Studies. 4 (2): 73–81. (2016).

57. Eastwood G, Cunningham AA, Kramer LD, Goodman SJ. The vector ecology of introduced *Culex quinquefasciatus* populations, and implications for future risk of West Nile virus emergence in the Galápagos archipelago. Medical Vet Entomology. 33 (1):44–55. (2019).

58. Hancock C, Camp JV. Habitat-specific host selection patterns of *Culex quinquefasciatus* and *Culex nigripalpus* in Florida. Journal of the American Mosquito Control Association. 38 (2):83–91. (2022).

59. Luna EC, Luna IS, Scotti L, Monteiro AFM, Scotti MT, De Moura RO, et al. Active essential oils and their components in use against neglected diseases and arboviruses. Oxidative Medicine and Cellular Longevity. 2019:1–52. (2019).

60. Huong LT, Hung NH, Linh NN, Pham TV, Dai DN, Hop NQ, et al. Essential oils of five *syzygium* species growing wild in Vietnam: chemical compositions and antimicrobial and mosquito larvicidal potentials. Molecules. 28 (22):7505. (2023).

61. Kek R, Hapuarachchi HC, Chung CY, Humaidi MB, Razak MABA, Chiang S, et al. Feeding Host Range of *Aedes albopictus* (Diptera: Culicidae) Demonstrates Its opportunistic host-seeking behavior in rural Singapore. Journal of Medical Entomology. 51 (4):880–4. (2014).

62. Li Y, Kamara F, Zhou G, Puthiyakunnon S, Li C, Liu Y, et al. Urbanization increases *Aedes albopictus* larval habitats and accelerates mosquito development and survivorship. PLoS Neglected Tropical Diseases. 8 (11):e3301. (2014).

63. Rose NH, Sylla M, Badolo A, Lutomiah J, Ayala D, Aribodor OB, et al. Climate and urbanization drive mosquito preference for humans. Current Biology. 30 (18):3570–3579.e6. (2020).

64. Fikrig K, Harrington LC. Understanding and interpreting mosquito blood feeding studies: the case of *Aedes albopictus*. Trends in Parasitology. 37 (11):959–75. (2021).

65. Fikrig K, Rose N, Burkett-Cadena N, Kamgang B, et. al. *Aedes albopictus* host odor preference does not drive observed variation in feeding patterns across field populations. Scientific Reports. 13 130. (2023).

66. Chen XG, Jiang X, Gu J, Xu M, Wu Y, Deng Y, et al. Genome sequence of the Asian Tiger mosquito, *Aedes albopictus*, reveals insights into its biology, genetics, and evolution. Proceedings of the National Academy of Sciences of USA. 112 (44). (2015).

67. Rose NH, Badolo A, Sylla M, Akorli J, Otoo S, Gloria-Soria A, et al. Dating the origin and spread of specialization on human hosts in *Aedes aegypti* mosquitoes. eLife. 12:e83524. (2023).

68. Zadra N, Rizzoli A, Rota-Stabelli O. Chronological incongruences between mitochondrial and nuclear phylogenies of *Aedes* mosquitoes. Life. 11 (3):181. (2021).

69. Hallem EA, Nicole Fox A, Zwiebel LJ, Carlson JR. Mosquito receptor for human-sweat odorant. Nature. 427 (6971):212–3. (2004).

70. Busula AO, Takken W, De Boer JG, Mukabana WR, Verhulst NO. Variation in host preferences of malaria mosquitoes is mediated by skin bacterial volatiles. Medical Vet Entomology. 31 (3):320–6. (2017).

71. Haertl T, Owsienko D, Schwinn L, Hirsch C, Eskofier BM, Lang R, et al. Exploring the interrelationship between the skin microbiome and skin volatiles: a pilot study. Frontiers in Ecology and Evolution. 11:1107463. (2023).

72. Thody AJ, Shuster S. Control and function of sebaceous glands. Physiological Reviews. 69 (2):383–416. (1989).

73. Oh J, Byrd AL, Deming C, Conlan S, Kong HH, Segre JA. Biogeography and individuality shape function in the human skin metagenome. Nature. 514 (7520):59–64. (2014).

74. Baker LB. Physiology of sweat gland function: the roles of sweating and sweat composition in human health. Temperature. 6 (3):211–59. (2019).

75. Kabelitz D, Schroder J. Mechanisms of epithelial defense (chemical immunology and allergy). 86:22–41 (2005).

76. Page MG, Heim J. Prospects for the next anti-*Pseudomonas* drug. Current Opinion in Pharmacology. 9 (5):558–65. (2009).

77. Buivydas A, Pasanen T, Senčilo A, Daugelavičius R, Vaara M, Bamford DH. Clinical isolates of *Pseudomonas aeruginosa* from superficial skin infections have different physiological patterns. FEMS Microbiology Letters. 343 (2):183–9. (2013).

78. Gadelhak GG, El-Tarabily KA, Al-Kaabi FK. Insect control using chitinolytic soil actinomycetes as bio-control agents. International Journal of Agriculture and Biology. 7 (4):627–633. (2005).

79. Dhanasekaran D, Sakthi V, Thajuddin N, Panneerselvam A. Preliminary evaluation of *Anopheles* mosquito larvicidal efficacy of mangrove actinobacteria. International Journal of Applied Biology and Pharmaceutical Technology. 1 (2): 374–381. (2010).

80. Devi Salam M. Antimicrobial potential of actinomycetes isolated from soil samples of Punjab, India. Journal of Microbiology and Experimentation. 1 (2): 63–68. (2014).

81. Fujii T, Shinozaki J, Kajiura T, Iwasaki K, Fudou R. A newly discovered *Anaerococcus* strain responsible for axillary odor and a new axillary odor inhibitor, pentagalloyl glucose. FEMS Microbiology Ecology. 89 (1):198–207. (2014).

82. Raji JI, DeGennaro M. Genetic analysis of mosquito detection of humans. Current Opinion in Insect Science. 20:34–8. (2017).

83. Bohbot JD, Dickens JC. Insect repellents: modulators of mosquito odorant receptor activity. PLoS ONE. 5 (8):e12138. (2010).

84. Greppi C, Laursen WJ, Budelli G, Chang EC, Daniels AM, Van Giesen L, et al. Mosquito heat seeking is driven by an ancestral cooling receptor. Science. 367 (6478):681–4. (2020).

85. CDC. Mosquitoes. 2024 [cited 2025 Oct 31]. About Mosquitoes. Available from: https://www.cdc.gov/mosquitoes/about/index.html

86. Liu F, Coutinho-Abreu IV, Raban R, Nguyen TTD, Dimas AR, Merriman JA, et al. Engineered skin microbiome reduces mosquito attraction to mice. PNAS Nexus. 3 (7):267. (2024).

87. Lommen A, Kools HJ. MetAlign 3.0: performance enhancement by efficient use of advances in computer hardware. Metabolomics. 8 (4):719–26. (2012).

88. Lommen A. MetAlign: Interface-driven, versatile metabolomics tool for hyphenated full-scan mass spectrometry data preprocessing. Analytical Chemistry. 81 (8):3079–86. (2009).

89. Tikunov YM, Laptenok S, Hall RD, Bovy A, De Vos RCH. MSClust: a tool for unsupervised mass spectra extraction of chromatography-mass spectrometry ion-wise aligned data. Metabolomics. 8 (4):714–8. (2012).

90. Callahan BJ, Wong J, Heiner C, Oh S, Theriot CM, Gulati AS, et al. High-throughput amplicon sequencing of the full-length 16S rRNA gene with single-nucleotide resolution. Nucleic Acids Research. 47 (18):e103–e103. (2019).

91. Bolyen E, Rideout JR, Dillon MR, Bokulich NA, Abnet CC, Al-Ghalith GA, et al. Reproducible, interactive, scalable and extensible microbiome data science using QIIME 2. Nature Biotechnology. 37 (8):852–7. (2019).

92. Rognes T, Flouri T, Nichols B, Quince C, Mahé F. VSEARCH: a versatile open source tool for metagenomics. PeerJ. 4:e2584. (2016).

93. Nearing JT, Douglas GM, Comeau AM, Langille MGI. Denoising the denoisers: an independent evaluation of microbiome sequence error-correction approaches. PeerJ. 6:e5364. (2018).

94. Walker SP, Barrett M, Hogan G, Flores Bueso Y, Claesson MJ, Tangney M. Non-specific amplification of human DNA is a major challenge for 16S rRNA gene sequence analysis. Scientific Reports. 10 (1):16356. (2020).

95. Zaccaron AZ, Stergiopoulos I. Characterization of the mitochondrial genomes of three powdery mildew pathogens reveals remarkable variation in size and nucleotide composition. Microbial Genomics. 7 (12). (2021).

96. Deissová T, Zapletalová M, Kunovský L, Kroupa R, Grolich T, Kala Z, et al. 16S rRNA gene primer choice impacts off-target amplification in human gastrointestinal tract biopsies and microbiome profiling. Scientific Reports. 13 (1):12577. (2023).

97. Dhariwal A, Chong J, Habib S, King IL, Agellon LB, Xia J. MicrobiomeAnalyst: a web-based tool for comprehensive statistical, visual and meta-analysis of microbiome data. Nucleic Acids Research. 45 (W1):W180–8. (2017).

